# Timing polymerase pausing with TV-PRO-seq: dissecting the interplay of pausing duration and location, and gene expression

**DOI:** 10.1101/461442

**Authors:** Jie Zhang, Massimo Cavallaro, Daniel Hebenstreit

## Abstract

Transcription of many genes in metazoans is subject to polymerase pausing, which is the transient stop of transcriptionally-engaged polymerases. This is known to mainly occur in promoter proximal regions but it is not well understood. In particular, a genome-wide measurement of pausing times at high resolution has been lacking. We present here the time-variant precision nuclear run-on and sequencing (TV-PRO-seq) assay, an extension of the standard PRO-seq that allows us to estimate genome-wide pausing times at single-base resolution. Its application to human cells demonstrates that, proximal to promoters, polymerases pause more frequently but for shorter times than in other genomic regions. Pausing release by the detergent sarkosyl, previously believed to be linked to the factor NELF at the promoter proximal region only, is independent of the latter. Comparison with single-cell gene expression data reveals that the polymerase pausing times are longer in highly expressed genes, while transcriptionally noisier genes have higher pausing frequencies and slightly longer pausing times. Analyses of histone modifications suggest that the marker H3K36me3 is related to the polymerase pausing.

## Introduction

RNA polymerase II (Pol II) is not moving uniformly after transcription initiation at metazoan genes; it frequently pauses during transcription (1–5). The most conspicuous phenomenon that has been extensively studied in this context is pausing at promoter proximal regions (PPRs) (6–8). Several protein factors such as negative elongation factor (NELF), (9) and DRB sensitivity-inducing factor (DSIF, (10) have been found to influence pausing, along with more generic factors, such as DNA sequence (11) and/or nucleosomes (6, 12). Another factor, positive transcription elongation factor-b (P-TEFb), mediates phosphorylation of NELF and DSIF, and facilitates Pol II release from promoter proximal pausing into active elongation (13, 14).

While polymerase pausing was discovered decades ago (15–17), its purpose remains uncertain. Several examples suggest a role in expression regulation, in particular for genes that need to respond quickly, as upon heat shocks, for instance (18). On the other hand, pausing appears to be common; it was reported to occur at roughly a third of all genes (2), as also demonstrated by small-molecule inhibition (with Flavopiridol, FP) of P-TEFb, which leads to a widespread suppression of all active genes (4, 13, 19). These results point towards a fundamental function of pausing in the transcriptional machinery. On the other hand, recent research suggests that high promoter proximal Pol II densities, which are usually interpreted as signatures of pausing, rather reflect a high turnover rate of nascent transcripts, i.e., abortive transcription (20–22). In fact, in vivo experiments show that only less than 10% of polymerases can escape from the promoter proximal region (PPR) and enter productive elongation (21).

Understanding of polymerase pausing has been greatly advanced by several types of assays based on next generation sequencing; these include ChIP-seq and ChIP-nexus (both based on chromatin immunoprecipitation) (23), GRO-seq (Global nuclear Run-On sequencing, (6), (m)NET-seq (mammalian Native Elongating Transcript sequencing) (24) and PRO-seq (Precision nuclear Run-On sequencing), (8), along with more recent developments, such as CoPRO-seq (Coordinated PRO-seq), which can correlate pausing with transcriptional start sites (25) and 5’ capping states (26). These assays are mostly based on the sequencing of polymerase-associated DNA fragments or nascent mRNA. After mapping the resulting sequencing reads to the genome, locations with higher read counts (‘peaks’) are thought to reflect greater polymerase occupancies, which are then used as proxies for pausing locations. Aside from revealing Pol II accumulations of polymerase in the PPR, these assays led to many other important findings (3, 26), including pausing sites at gene body(7) and 3’ ends of genes (6).

A fundamental problem with these methods based on measuring polymerase occupancy is that they cannot separate the influence of pausing and the turnover of aborted transcripts in the promoter proximal region. This is due to the methods’ inability to discriminate between few slow polymerases and many fast polymerases detected at a genomic position during a given amount of time; both cases will result in identical peaks of sequencing reads, which prevents measuring the actual pausing times. The latter has been accomplished only via blocking transcription initiation with the covalent TFIIH subunit XPB inhibitor Triptolide (27) and measuring Pol II release dynamics from the promoter proximal region) and at low resolution (28, 29); to the best of our knowledge, genome-wide data for pausing times at single positions are lacking.

We present here an extension of the PRO-seq assay, which we termed time-variant PRO-seq (TV-PRO-seq), that achieves this goal. TV-PRO-seq removes the influence of many factors that affect Pol II occupancy (such as expression level, abortive transcription and pausing fraction, see Box 1) thus allowing us to directly study the pausing times in vitro. The TV-PRO-seq results are consistent with the more limited data obtained from in vivo Trp treatment followed by sequencing, and go beyond the latter in revealing, based on genome wide analyses, the novel finding that Pol II pauses more often but for shorter times at each base in the PPRs than in other regions of a gene. The pausing related to NELF can be lifted by sarkosyl treatment. Our results also show that pausing within the genes with higher expression levels lasts longer. Genes with higher transcriptional noise tend to have higher NELF levels in their PPRs, along with higher densities of pausing sites that extend to their gene bodies yet differ little in terms of pausing times. Our analysis of individual pausing sites has also yielded insights into the pausing profiles associated with productive elongation; we find that the active elongation marker H3K36me3 surprisingly relates to pausing, suggesting pausing could establish H3K36 methylation and/or a dynamic equilibrium of H3K36me3 and histone acetylation that contributes to elongation rate regulation.

### Box 1. Schematic illustration of pausing-related phenomena and terms.

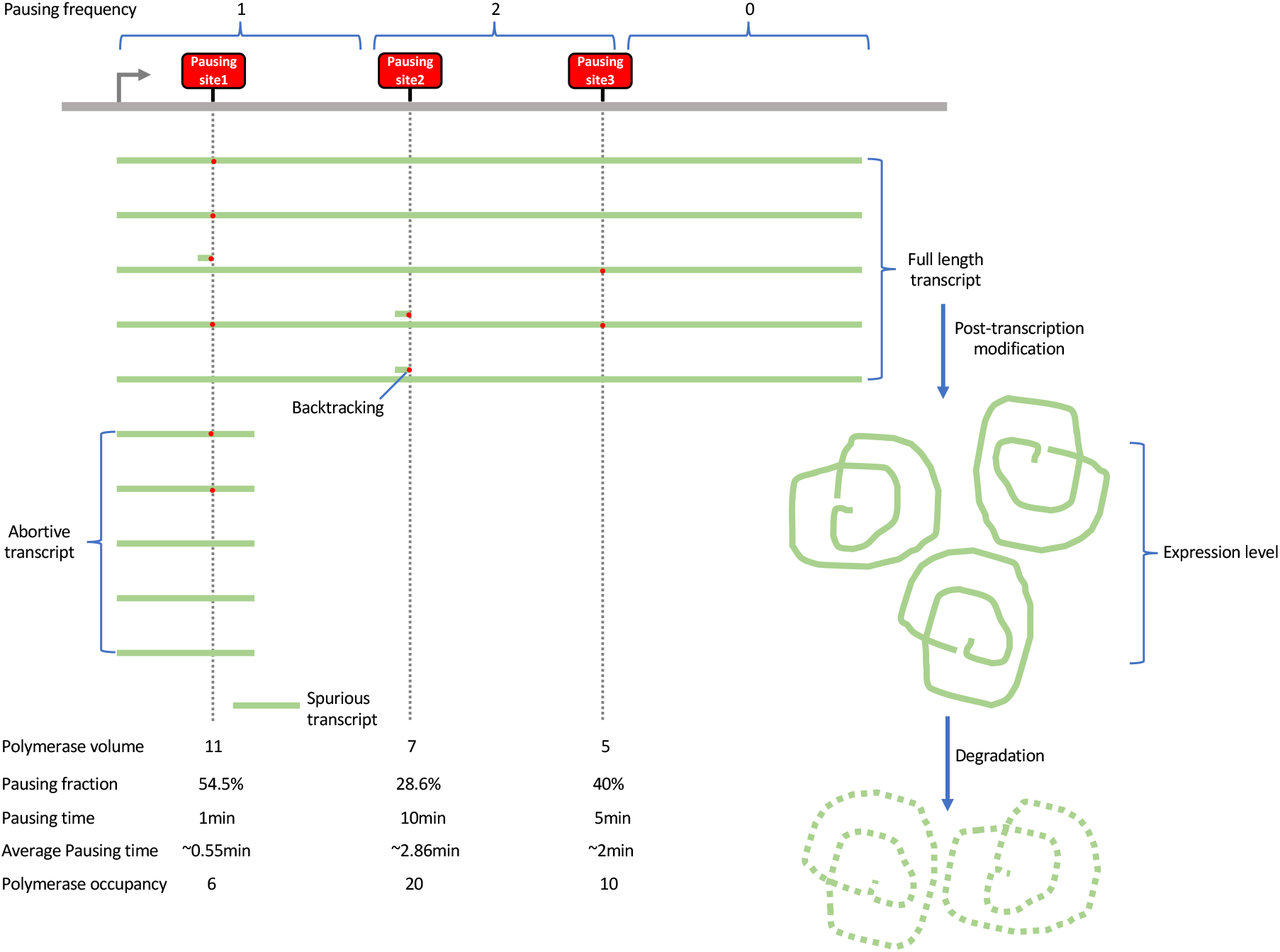

## Results

### Estimating pausing times for individual pausing sites in genome-wide fashion

Numerous advanced sequencing methods have been developed for studying pausing based on RNA polymerase occupancy (6, 8, 23, 24). However, RNA polymerase occupancy correlates with various parameters, such as gene expression level, polymerase turnover rate, pausing fraction and pausing time, thus preventing independent measurement of the latter (Box 1). To overcome this problem, we developed TV-PRO-seq; it can extract pausing times of polymerases at individual pausing sites in genome-wide fashion. This enables us to dissect pausing profiles to gain a mechanistic understanding.

We developed TV-PRO-seq based on a detailed analysis of the principles underlying PRO-seq. The assay relies on the replacement of native NTPs in the nuclei with biotin-labelled ones (biotin-NTPs), which become incorporated into the 3’ end of nascent RNA (30) over a short period of time (‘run-on’ time). This blocks further transcription and makes polymerase drop off the template, thus marking the exact location of incorporation. The biotin tag is then used to isolate newly synthesized RNA, followed by library preparation and sequencing. The longer the run-on time, the more polymerases will be released. Eventually, all polymerases will have been released and no more reads can result. Although the kinetics of biotin-NTP incorporation are not necessarily identical to those of physiological NTP, we argue that they are correlated in rank order to those in-vivo, thus permitting inference of biological dynamics. Preparing several PRO-seq reactions using different run-on times allows us to fit saturation curves, whose slopes permit estimation of pausing (-release) times (Fig. 1A).

**Fig. 1.**
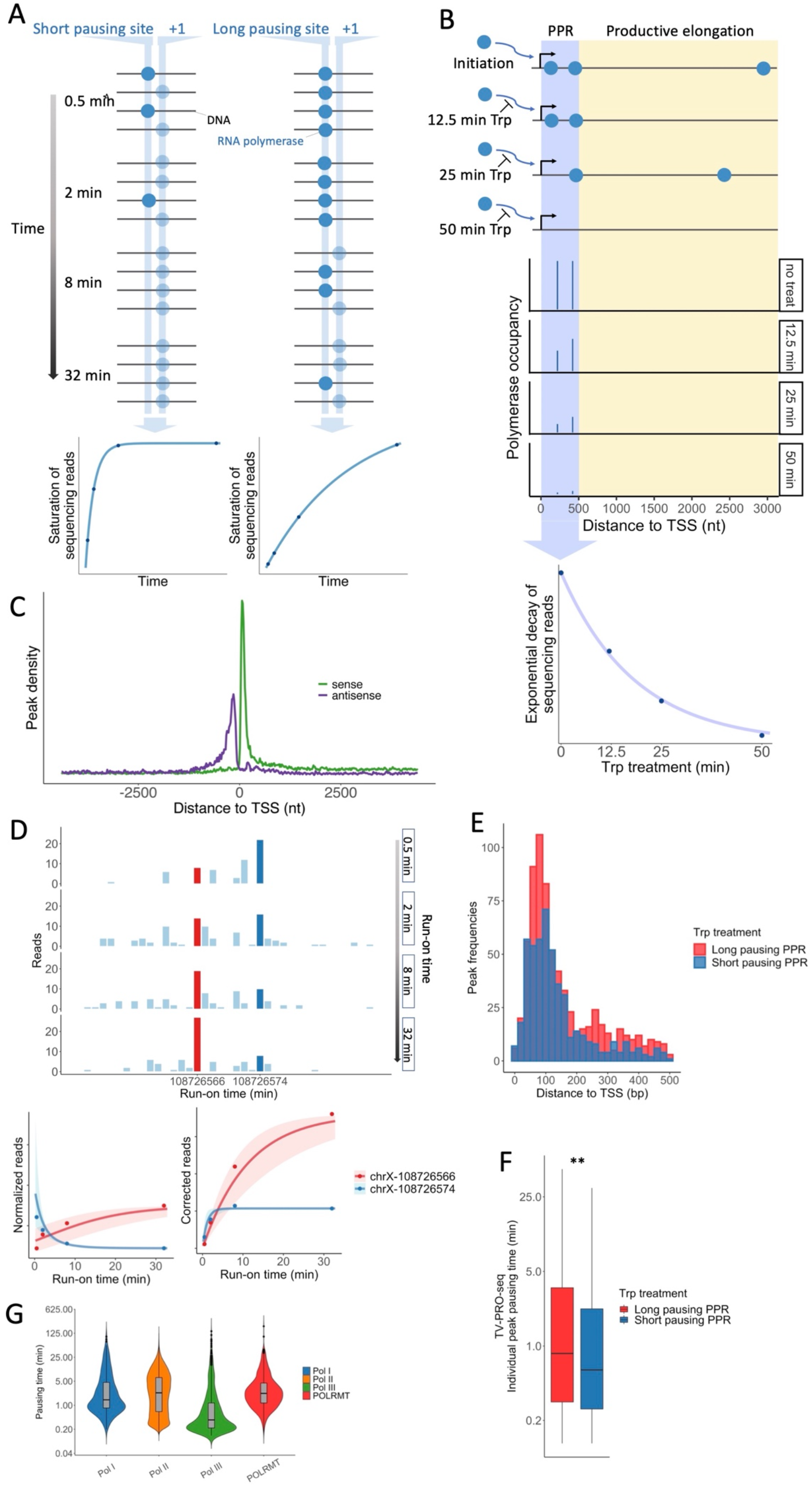
Principle of TV-PRO-seq. (**A**) The black horizontal lines symbolize a generic DNA region with a short (left graphics) and a long (right graphics) pausing site. The blue dots symbolize RNA polymerases that either are stationary or have just moved by one position (and incorporated a biotinylated NTP) as indicated by the lighter blue shades. A sequencing read results at a position if a polymerase steps forward by one base. Eventually, all polymerases will have moved, i.e., all positions will be saturated. The saturation takes longer at the position (‘+1’) adjacent to the long pausing site, since the polymerases are released at a lower rate than from the short pausing site. Saturation curves (lower plots) can be inferred by reads from a run-on time-course at each position, genome-wide. (**B**) Trp blocks transcription initiation, thus decreasing the polymerase occupancies at the PPRs. The decay rate at different pausing sites is also influenced by their distance to TSS. Two pausing sites with the same pausing times are represented in the diagram; the decay rate of polymerase occupancy of the most downstream peak is underestimated by the presence of persisting polymerases upstream. The total reads of the PPR from Trp treatment based sequencing can be used to estimate the average pausing time in the PPR by fitting an exponential decay curve. (**C**) Distributions of sense and antisense read around TSSs from pooled TV-PRO-seq samples confirm high library quality. (**D**) Read numbers from two neighbouring peaks (red and blue bars) in chromosome X obtained at the different run-on times (top panel). Normalising these by the total-genome reads permits parameter estimation and produces the curves in the left bottom panel. Correcting by the total-genome read trend reveals the saturation curves in the right bottom panel (details in Suppl. Methods). Shaded regions are interquartile posterior ranges. (**E**) More peaks are found in the ‘long pausing’ PPR. The 2,000 genes with the highest polymerase occupancy in the PPR (first 500bp downstream of TSS) were used for analysis. 500 genes each retaining the highest and lowest polymerase occupancies after 10 min Trp treatment were grouped as ‘long pausing PPR’, and ‘short pausing PPR’, respectively. 702 peaks were identified in the long pausing PPR and 493 peaks were found in the short pausing PPR (Exact Binomial Test, P<10^-8^). (**F**) Peaks in the long pausing PPR have longer pausing times as measured by TV-PRO-seq (Mann-Whitney U test, P < 0.01). Peaks were grouped same as in (E). (**G**) Distributions of estimated pausing times for peaks in loci transcribed by Pol I, II, III and POLRMT. For all pairwise comparisons except Pol II vs POLRMT and Pol II vs Pol I (n.s.), P < 0.01, Bonferroni-corrected Mann-Whitney U test.

In TV-PRO-seq, polymerases theoretically can only move a maximum of one nucleotide after release from pausing during run-on (Fig. 1A). This allows us to record data for each nucleotide thus enabling genome-wide estimation of pausing times at single nucleotide resolution. The most widely used sequencing method for estimating pausing times, Triptolide (Trp) treatment based sequencing, is incapable of achieving this (28, 29, 31, 32). Trp is a covalent inhibitor of the TFIIH subunit XPB (33) and therefore blocks transcription initiation. Fitting decay curves to the (declining) polymerase occupancy of the region downstream of TSS upon a Trp treatment time series allows estimation of the average pausing times at the PPRs of all genes (Fig. 1B). As it prevents initiation, this method can only measure the average residence times (Box 1) in a small region downstream of the TSS; its application in the gene body or to individual pausing sites in the PPR is not possible, which is a significant limitation.

We performed TV-PRO-seq for HEK293 cells using 0.5, 2, 8, and 32-min run-on times.

The standard PRO-seq preparation buffer contains sarkosyl, an anionic detergent that has been reported to facilitate pausing release especially in the PPR of coding genes and enhancers (34, 35). To remove its effects towards pausing, we excluded sarkosyl from the run-on buffer of all TV-PRO-seq samples, except for an initial comparison.

To analyse the resulting data, we first called peaks based on a heuristic thresholding (Supp. Methods, Fig. S1), which resulted in 66,089 individual peaks for the HEK293 data. Plotting the distribution of peaks around TSSs reproduced the familiar pattern of promoter-proximal peaks and divergent transcription on the other strand (Fig. 1C). A replicate, with 47,713 peaks detected, shows a similar pattern of peak distribution (Fig. S2A), and the two replicates’ pausing times were consistent among them (Fig. S2B).

For comparison, we also generated two replicates of TV-PRO-seq with sarkosyl (Fig. S2C). Due to the lack of elongation rate measurements under sarkosyl treatment and the latter’s effects on pausing in general, we do not expect our estimates for the sarkosyl run-on samples to reflect the real pausing times (28, 36). The pausing times of these sarkosyl run-on samples have better consistency compared to sarkosyl-free samples (Fig. S2B, C), as the sarkosyl increases the run-on efficiency (35). In contrast, a complex pattern emerges when comparing sarkosyl and sarkosyl-free samples, indicating substantially stronger treatment effects than replicate variation in our datasets and confirming their informative value (Fig. S2D).

We constructed a mathematical model that takes account of our theoretical considerations; the model predicts the saturation curve as a function of the pausing release rate and the TV-PRO-seq run-on time (Supp. Methods). Fitting our saturation model to the time-course data of a set of peaks allows the inference of their pausing release rates, whose reciprocals are the pausing times. The model is embedded into a Bayesian framework, detailed in Supp. Methods. Examples of fitted curves corresponding to two close individual peaks are shown in Fig. 1D (saturation curves from the replicate data for the same peaks are shown in Fig. S3). Note that the saturations are subject to trade-offs in terms of sequencing reads with peaks with extremely high pausing times (whose polymerase occupation remains virtually unchanged during the time course experiment): normalizing the samples’ total read numbers to stay constant throughout the run-on time, the latter’s sizes will decrease (Fig. 1D, bottom left; see Methods for details). Our estimates of the individual pausing times are based on average elongation rates from published data (28, 37), which were incorporated into the model as a Bayesian informative prior (Supp. Methods).

Trp treatment followed by sequencing has been widely used for estimation of pausing times of PPR regions(28, 29, 31, 32). The quicker the reduction of polymerase occupancy upon after Trp treatment, the shorter the pausing times are assumed to be (Figure 1B). As an *in vitro* method, TV-PRO-seq shows a good consistency with Trp treatment. We took the 2,000 genes with the highest nascent transcription signal in PPR region. Within these, we identified the 500 genes with the most reduction of polymerase occupancy after 10 min Trp treatment as ‘short pausing PPR’, and the 500 with the least reduction as ‘long pausing PPR’. We then looked at the pausing times of peaks we identified by TV-PRO-seq within the two groups of PPRs. It shows that the genes with overall longer pausing in the PPR (Trp treatment following sequencing) have both more pausing sites (TV-PRO-seq, Figure 1E) and longer pausing time for each peak (TV-PRO-seq, Figure 1F).

The positions losing polymerase quicker after 10min Trp treatment only have short average pausing times according to TV-PRO-seq, which amount to less than half of those with high polymerase occupancy after Trp treatment (Fig. S4A). As the Trp acts from the TSS, the positions far from TSS would lose polymerase slower than the upstream ones (Figure 1B). This bias does not occur in TV-PRO-seq as the interruption of transcription happens at the position the polymerase is located at rather than the TSS. Thus we zoomed in to pausing sites within the first 500bp or even 100bp downstream of TSS to reduce this bias, and the difference remains significant (S4B and C).

Pol I and Pol III transcription contribute about 80% of the total RNA production in rapidly growing cells (38). Many noncoding RNAs that are essential for cells are transcribed by Pol I and Pol III. Some of these noncoding RNAs, such as RNase P, RNase MRP or transfer RNAs (tRNAs) are only about hundred nucleotides long (38–40). Previous methods for estimating pausing levels cannot work for these short genes, as they are mostly based on the so-called ‘pausing index’ (PI, also referred to as stalling index or escaping index, Supp. Methods), which normalizes polymerase occupancy within the first ~ 200 to 300nt downstream of TSSs by that in the gene body. The purpose of this normalization is to correct for the gene’s expression level by treating promoter proximal pausing as the (increased) relative occupancy over the gene body’s (6, 7). In contrast, TV-PRO-seq estimates pausing times for individual pausing sites, thus providing a way to study pausing in both short and long genes, and at any position. We pooled the pausing positions in Pol III transcribed genes and compared their times with those of pausing sites related to Pol I, Pol II and Mitochondrial polymerase (POLRMT). We found that pausing time varies for different type of polymerases. The long pausing times are mostly associated with Pol II, and surprisingly, Pol III pauses for the shortest times, overall (Fig. 1G).

We next had a closer look at tRNA genes in order to explore genic pausing time patterns. The clear pattern of pausing at tRNA genes demonstrates the power and precision of TV-PRO-seq to reveal significant differences of pausing time at single base resolution (Fig. S5A, Fig. S5B). Abundant short intragenic pausing concentrates in three peaks that appear conserved across genes. Interestingly, the longest pausing times are specifically found at the transcription end sites (TESs). A region downstream of TESs also shows an enrichment of pausing sites and has slightly longer pausing times compared to the gene body. Releases from both intragenic and TES-related pausing sites are triggered by sarkosyl (Fig. S5C). Overall, pausing times at TESs and the adjacent downstream regions are still longer than the intragenic ones under sarkosyl-including run-on conditions.

### TV-PRO-seq estimates pausing times independently of confounding factors

Pol II is found to be enriched in promoter proximal regions (6, 41), which is usually interpreted as longer Pol II residence time and thus pausing in these regions than elsewhere (Fig. 2A, B). This pausing is often claimed to be of particularly long duration, from 2mins to even 30mins for some genes, based on studies blocking transcription initiation with Trp and the slow reduction of polymerase occupancy that ensues (28, 29, 31, 32). However, measuring pausing time in this way relies on a rapid uptake of Trp. Yet, 500nM Trp treatment, which is the usual concentration used for initiation inhibition prior to pausing time measurement in PPR regions(28, 29), has proved ineffective in this respect (42).

**Fig. 2.**
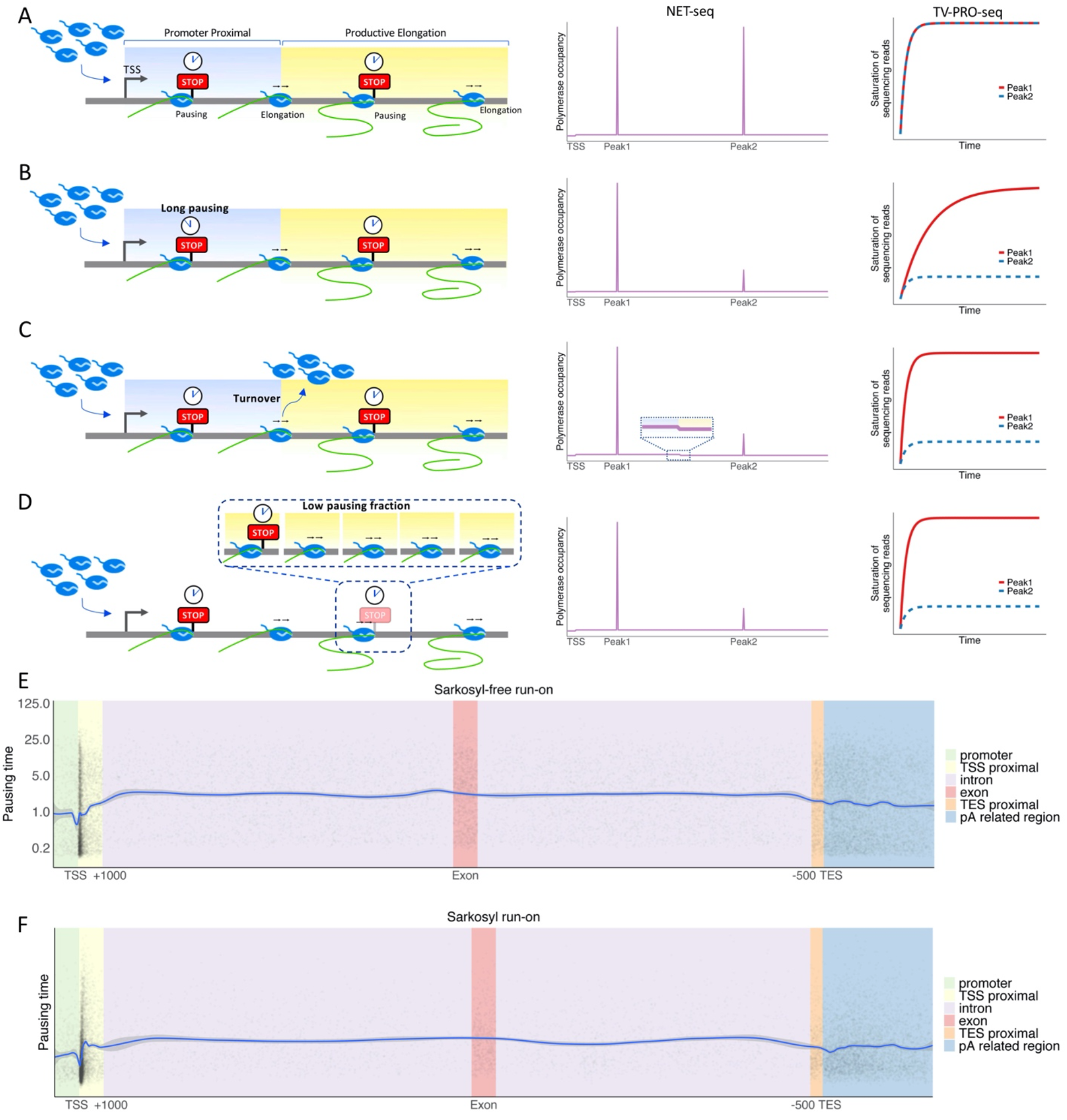
Several factors influence polymerase occupancy. (**A**) The left panel shows a schematic example case by in silico simulations; two regions were designated as ‘promoter proximal’ (blue shading) and ‘productive elongation’ (yellow shading), each with a single pausing site (Peak1 and Peak2) with identical properties. Polymerase occupancies measured by NET-seq (middle panel) and the saturation curves resulting from TV-PRO-seq (right panel) of the two peaks will be the same. (**B**) As (A), but the pausing time of Peak1 was set five times longer (clock symbols) than Peak2’s. Both, polymerase occupancy and pausing time (1/β_i_ see Supp. Methods) of Peak1 would be measured to be five times higher than Peak2’s. (**C**) As (A) and (B), but 80% of polymerase is assumed to abort transcription at the boundary of the promoter proximal region thus reducing by 80% polymerase occupancy in the productive elongation region. Therefore, the measured polymerase occupancy of Peak1 would still be 5-fold higher than at Peak2, for both, NET-seq and TV-PRO-seq. However, in contrast to NET-seq, TV-PRO-seq is still able to correctly measure the pausing times at the two peaks to be equal despite their differing sizes. In contrast to (B, D), high abortive transcription would also decrease the polymerase occupancy in the productive elongation region (magnified section). (D) As (A), but only one fifth of the polymerase is assumed to pause at Peak2 (i.e., the pausing fraction is a fifth), thus its polymerase occupancy would decrease to 1/5 of Peak1’s. The pausing time of Peak2, however, would be about the same as Peak1’s. (**E**) Pausing times at mRNA-transcribing metagene. Each grey dot represents a pausing peak, with corresponding pausing time given by its y-axis value. The x-axis values correspond to the absolute position within −/+ 1000 nt of the TSS (green and yellow tinged regions, respectively). The intron/exon regions (purple/red, respectively) start after +1000nt of the TSS and end before −500 of the TES (introns were split into an upstream and a downstream group at the gene’s middle point) and 500nt upstream and 4500nt downstream of the TES were indicated (orange and blue, respectively). The blue line corresponds to the moving average (LOESS fit). The grey shading indicates the 0.95 confidence interval and is negligible on this scale, hence invisible over most of the graph. The widths of exons and introns have been scaled to their relative average lengths. (**F**) Similar to (E), but including sarkosyl during the run-on reactions.

Indeed, polymerase densities higher in the PPR than in the regions downstream can have different causes. If transcription aborts before entering productive elongation, the polymerase occupancy will appear higher in the PPR (Fig. 2A, C); in fact, recent research shows that Pol II does have a high turnover rate in promoter proximal regions (20–22). Modelling based on in-vivo experiments suggests that only about 1/13 of Pol II can escape from the PPR and progress into productive elongation (21); in contrast to reported average pausing times from 7min to up to half an hour proximal to promoters (28, 29, 31), this in-vivo study claims polymerase residence times in the PPR of only about 42s (21). Similarly, a lower pausing fraction, i.e., lower utilization of pausing sites (Box 1) by polymerase engaged in productive elongation, yields lower occupancy downstream of the PPR (Fig. 2A, D). Also, a single pausing site with long pausing time vs many short pausing sites will result in the same contribution to the PPR’s overall polymerase occupancy.

Even though various sequencing methods have been developed/used for the research on transcriptional dynamics and similar topics (6, 8, 23, 24, 26), these are in fact restricted to reveal polymerase occupancy only. Extending the pausing time, increasing the turnover rate or boosting utilization of a pausing site can affect polymerase occupancy in similar ways (Fig. 2A-D, NET-seq). To distinguish the source of polymerase enrichment among these three possibilities, we developed TV-PRO-seq. In contrast to previous approaches using a time series of Trp treatment (Fig. 1B) followed by ChIP-seq or GRO-seq (28, 29, 31, 32), TV-PRO-seq can not only remove the influence of early-terminated transcripts, but also measure pausing times without Trp treatment. Both of these advantages prevent over-estimating pausing times in the PPR, since the artefactual extra occupancies due to aborted transcripts and slow Trp uptake do not influence the results.

As shown in Figure 2E, even though we find a much higher frequency of polymerase pausing in the PPR, TV-PRO-seq demonstrates that, in fact, individual pausing events close to TSSs last shorter times on average than those in other regions. Our interpretation of this is that pausing in the PPR is more akin to a collection of check points with high possibilities to pause polymerase for short times rather than a unitary long pausing apparatus which holds it back from moving into productive elongation. Even though individual pausing events in the PPR are shorter, considering the higher pausing frequency and potentially higher pausing fraction, the average elongation speed of polymerase in the PPR might still be lower than further downstream.

### Sarkosyl facilitates release from NELF-mediated long pausing

Sarkosyl specifically facilitates pausing release in the PPR (35). This effect is reflected in TV-PRO-seq by the deepening of the dip in pausing times downstream of TSSs (Fig. 2E,F and Fig. 3A). Also, more peaks in the PPR are found in the sarkosyl run-on sample compared to the sarkosyl-free one (Fig. 3B). This is due to PRO-seq’s selective detection of active polymerases; sarkosyl appears to denature NELF, thus causing a pausing release in the PPR and boosting peak density (35).

**Fig. 3.**
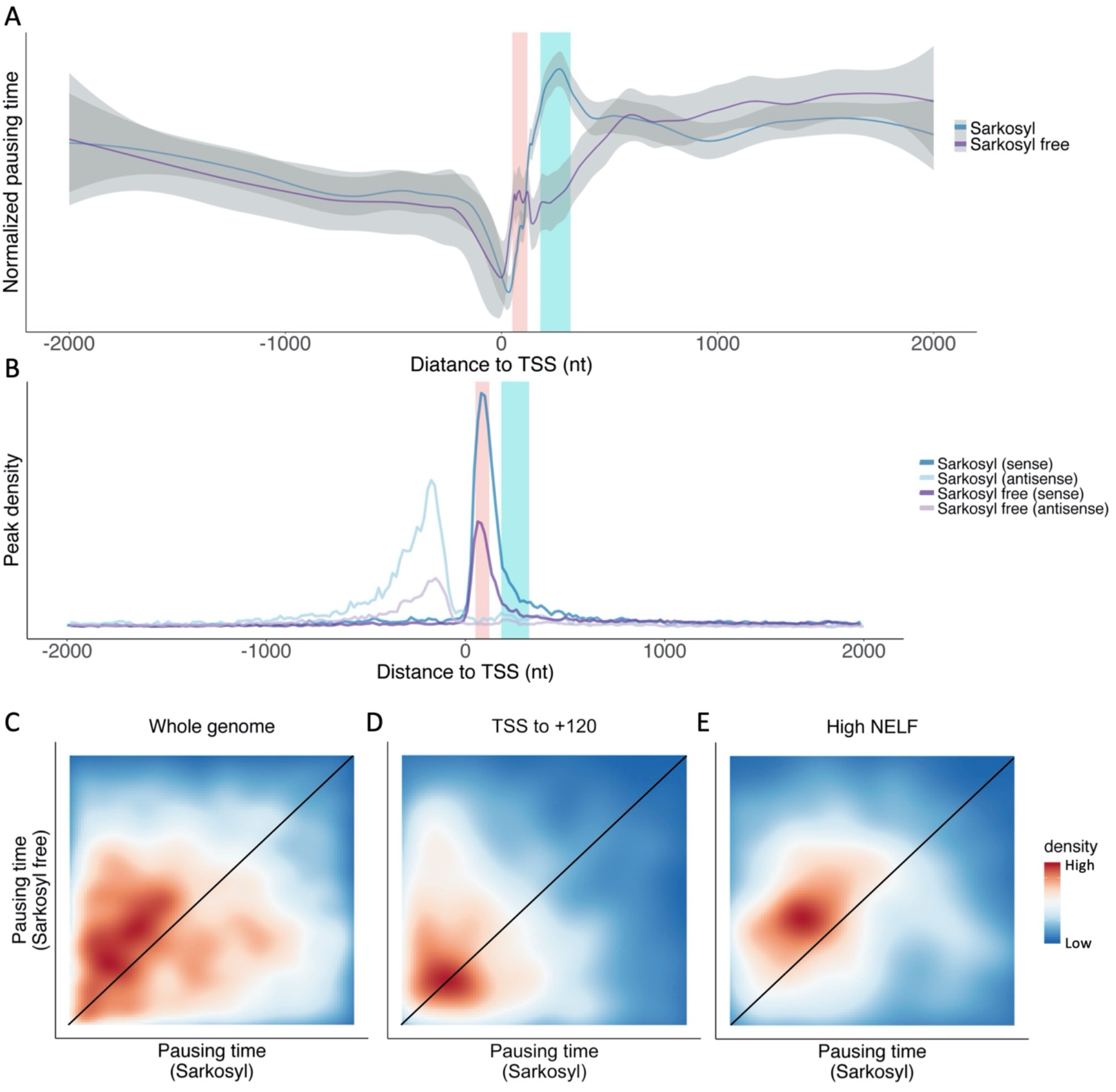
Influence of sarkosyl on pausing. (**A**) Pausing time of peaks around TSS. For removing the systematic bias of pausing time estimation the average pausing time is normalized to the same value for samples with sarkosyl (blue line) or with a sarkosyl-free (purple line) run-on. Pausing in the +50 to +120 region (pink shading) is sensitive to sarkosyl, while pausing in the +180 to +320 region (cyan shading) shows resistance to sarkosyl. (**B**) Sarkosyl increases the PPR peak density for both sense and divergent transcription. (**C**) 2D density plots show the pausing time rank of the equivalent peaks in sarkosyl sample and sarkosyl-free sample. The black line reflects peaks with intermediate influence on pausing time by sarkosyl. Peaks above the black line correspond to pausing sites releasing paused polymerase after sarkosyl treatment. (**D**) Similar to (C), only for the peaks within the first 120bp of genes. (**E**) Similar to (C), but only peaks with top 10% of NELF level.

NELF and DSIF are the best characterized factors involved in promoter proximal pausing (2, 43, 44) and their depletion significantly reduces polymerase occupancy in the promoter proximal region (12, 45). P-TEFb is a necessary factor for productive elongation which facilitates dissociation of NELF and converts DSIF to a positive elongation factor by phosphorylating it (2, 43). Inhibition of P-TEFb can prevent polymerase from entering productive elongation at nearly all active genes (4, 13, 19).

Interestingly, we found that the effect of pausing release caused by sarkosyl is mostly restricted to the first 120bp downstream of TSS (Fig. 3A), which coincides with the region with the highest NELF levels (Fig. S6A-B). The situation differs further downstream; polymerases show long pausing times between +180 to +320 (Fig. 3A). But when we plot the pausing time of peaks with different NELF coverage around TSSs of the sarkosyl-free sample (Fig. S6C), we find NELF to correlate with higher pausing times only within the first region, +120 downstream of TSS. In the region further downstream, pausing sites with higher NELF levels actually have shorter pausing times than the other peaks at the same distance towards the TSS (Fig. S6C).

To further dissect the relation between NELF and pausing, we revisit the sarkosyl-treated samples. Sarkosyl disturbs pausing in the PPR (Fig. S6D), but its effect on paused polymerase differs varies with distance to TSS. For the first 120bp downstream of TSS, all peaks with different NELF levels show a reduction of pausing time. Pausing with low NELF levels does not show a pausing time decrease within the +180 to +320 region, though. We suspect that this indicates that pausing is established by different mechanisms whose prevalence varies with the type of region. Candidates include pausing related to G-quadruplex DNA secondary structures, which, similar to NELF/DSIF, are also enriched in the PPRs (11). Nucleosomes also can pause polymerase, specifically at the +1 nucleosome (2, 43, 44), but also further downstream in the gene body (8, 12, 46, 47). Pause elements (48, 49) and nascent RNA structures (50) can induce pausing in the whole gene. In fact, peaks in the first 120bp were not affected by sarkosyl more than other peaks (Fig. 3C-D). In contrast, a strong effect can be seen for the peaks with high NELF level (Fig. 3E). Specifically, the peaks which most sensitive to sarkosyl are those peaks in gene body rather than PPR (Fig. S6E, F).

### Polymerase pausing and expression level

Counterintuitively, the presence of polymerase pausing is not associated with low gene expression. As a matter of fact, most pausing is found in active genes (2). In line with this, paused polymerases in the PPR have been suggested to keep the chromatin in an open state thus keeping the gene active (12). On the other hand, these paused polymerases have been described to block initiation of successive polymerases (29).

Studies of polymerase pausing and expression levels have been mostly focused on the PPR. With TV-PRO-seq, we can produce a more detailed picture across the whole genome. Highly expressed genes should have more detectable pausing sites due to higher polymerase volume. As expected, we find about 10 times more pausing sites in highly expressed genes (top 20% quantile; Fig. 4A). Interestingly, highly expressed genes do not only have more pausing sites, but also higher pausing times at each site (Fig. 4B). This difference is maintained across the whole gene, but is especially prominent in the PPR and downstream of TESs (Fig. 4C-D).

**Fig. 4.**
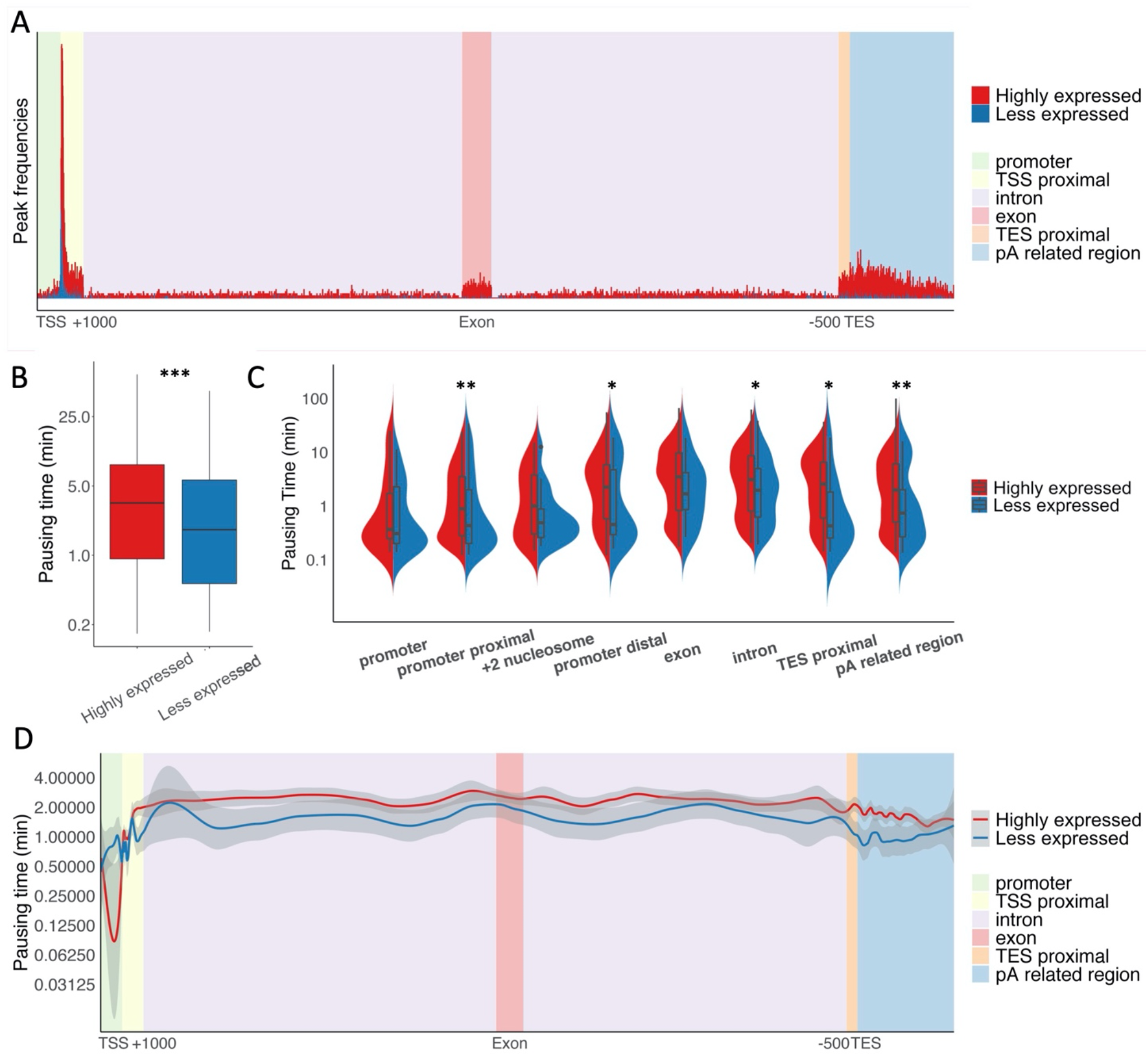
Pausing profiles and expression level. (**A**) Absolute peak density at mRNA-transcribing metagene as in Fig. 2E, for genes classified into different expression levels (‘highly expressed’, ‘less expressed’; red, blue, respectively). (**B**) Pausing times of pausing sites in highly expressed genes are longer than less expressed ones. P < 10^-23^, Mann-Whitney U test. (**C**) Pausing times of different regions of highly expressed and less expressed genes. Definitions of the region are the same as in Fig. 2E, TSS proximal have been split into promoter proximal (TSS to +120), +2 nucleosome (+180 to +320) and promoter distal (+500 to +1000) according to the different effects of sarkosyl on these regions. For promoter proximal and pA related region, P < 0.01; promoter distal, intron and TES proximal, P < 0.05; Mann-Whitney U test. (**D**) Pausing times of pausing peaks among genic regions for ‘low’ and ‘high’ expression genes at the metagene as in (A) shown as LOESS fits as in Fig. 2E.

Despite this, highly expressed genes do not appear to elongate more slowly (51). This may be due to biased sampling, since elongation rate measurements can only be taken for long genes. Conversely, TV-PRO-seq is not restricted in this way, which allowed us to revisit the pausing time analysis for peaks in extremely long genes and relatively short genes (>100kb & 3-10kb, respectively). Interestingly, we found that pausing in the extremely long genes tends to be shorter than in the short genes (Fig. S7A), the difference being significant only for highly expressed genes (Fig. S7B-C). This dilutes the difference in pausing times between highly expressed and less expressed genes among long genes (it results in a non-significant difference between these groups) albeit highly expressed long genes do have slightly higher pausing times than their less expressed counterparts (Fig. S7D).

We suggest that these different pausing profiles relate to the regulation of gene expression. The rate-limiting step of the less expressed genes is considered to be the activation/deactivation of the promoter. In highly expressed genes, instead, the promoter is thought to be always active, thus, the regulation of expression must, to a certain degree, rely on post-initiation mechanisms, including the pausing. Short genes might provide insufficient space in their gene bodies for the complex regulatory machinery to adjust expression and therefore denser and longer pausing might be utilized in these genes, instead.

### Polymerase pausing and transcriptional noise

A gene’s expression level is determined by its initiation rate, the fraction of nascent RNA that is turned into mature RNA, and the latter’s degradation rate. Polymerase pausing also influences the gene expression by adjusting the elongation process, but this chiefly results in the dispersed distribution of mRNAs among individual cells rather than contributing to the mean expression (52). This dispersion, or ‘noise’, is quantified by the CV^2^ (square of coefficient of variation) and can be obtained in genome-wide fashion from single-cell RNA-seq (e.g., Drop-seq) data. To study the relation between noise and pausing, we used Drop-seq data for HEK293 cells (53) and classified genes based on their CV^2^ for a moving average of mean expression levels. This reduces the influence of the latter, which the noise depends on (54, 55) (Fig. S8A).

We assigned genes to ‘low-’ and ‘high noise’ classes (Fig. S8B-C). We find that, overall, noisier genes have significantly higher pausing frequency (the number of pausing peaks in a given region, Box 1) throughout gene bodies (Fig. 5A). Polymerase also tends to pause longer in noisier genes, albeit a statistically significant difference emerges only for introns (Fig. 5B-D). Our TV-PRO-seq based analyses agree with previous theoretical considerations that predict more and longer pausing for high-noise genes (52, 56, 57). Overall, highly expressed genes with high transcriptional noise tend to have more and longer pausing than other genes. Our result suggests the importance of pausing in regulation of expression (Fig. S9).

**Fig. 5.**
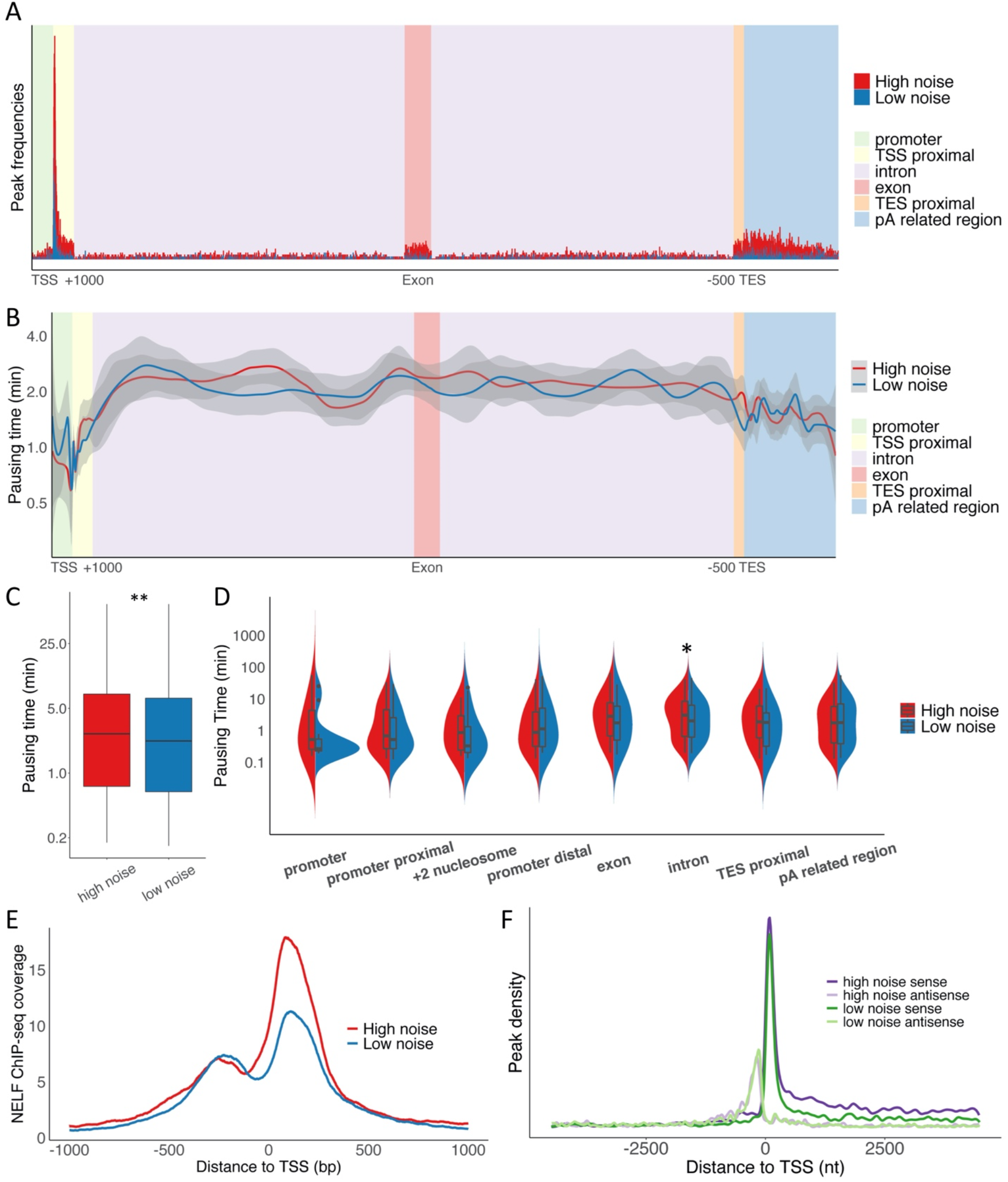
Pausing profiles and transcriptional noise. (**A**) Absolute peak density at mRNA-transcribing metagene as in Fig. 2E, for genes classified into different levels of transcriptional noise (‘high’, ‘low’; red, blue, respectively). (**B**) Pausing times of pausing peaks among genic regions for ‘low’ and ‘high’ noise genes at the metagene as in (A) shown as LOESS fits as in Fig. 2E. (**C**) Pausing times of pausing sites in high noise genes are longer than those of low noise genes. P < 0.01, Mann-Whitney U test. (**D**) Pausing times of different regions of high and low noise genes. Definitions of the region are the same as in Fig. 4C. For introns, P < 0.01, Mann-Whitney U test. (**E**) NELF coverage at TSSs of genes with high or low noise. (**F**) Absolute peak densities of both sense and antisense transcription of high or low noise genes.

We also find NELF levels to correlate with the noise of genes (Fig. 5E). For divergent transcription, NELF levels differ little between high and low noise genes (Fig. 5E). This can explain why the divergent transcription of genes with different noise levels has similar pausing frequencies in both (Fig. 5F).

### Histone modification and polymerase pausing

After these results, we turned our attention towards pausing and chromatin states. Different types of histone modifications influence transcription in various ways and vice versa (58). For instance, new histone acetylation is found at many genes after a heat shock (44, 59). Histone acetylation can also accelerate the release of paused polymerase (47, 60–62).

We thus investigated the effects of chromatin states on pausing times. To this end we classified peaks into ‘long’ and ‘short’ according to their pausing times and quantified their presence around different chromatin features. We found relations between pausing times and DNA accessibility and/or regulatory character; open chromatin regions as determined by DNase-seq display strong enrichment of short pausing (Fig. 6A). This is consistent with our other results, as open chromatin is found at the PPR, which in turn is enriched for NELF (Fig. S6A). The dramatic drop of DNA accessibility we see after the short pausing sites suggests that polymerases tend to pause in front of closed chromatin. A similar drop, albeit of reduced magnitude, is also seen for long pausing (Fig. 6A).

**Fig. 6.**
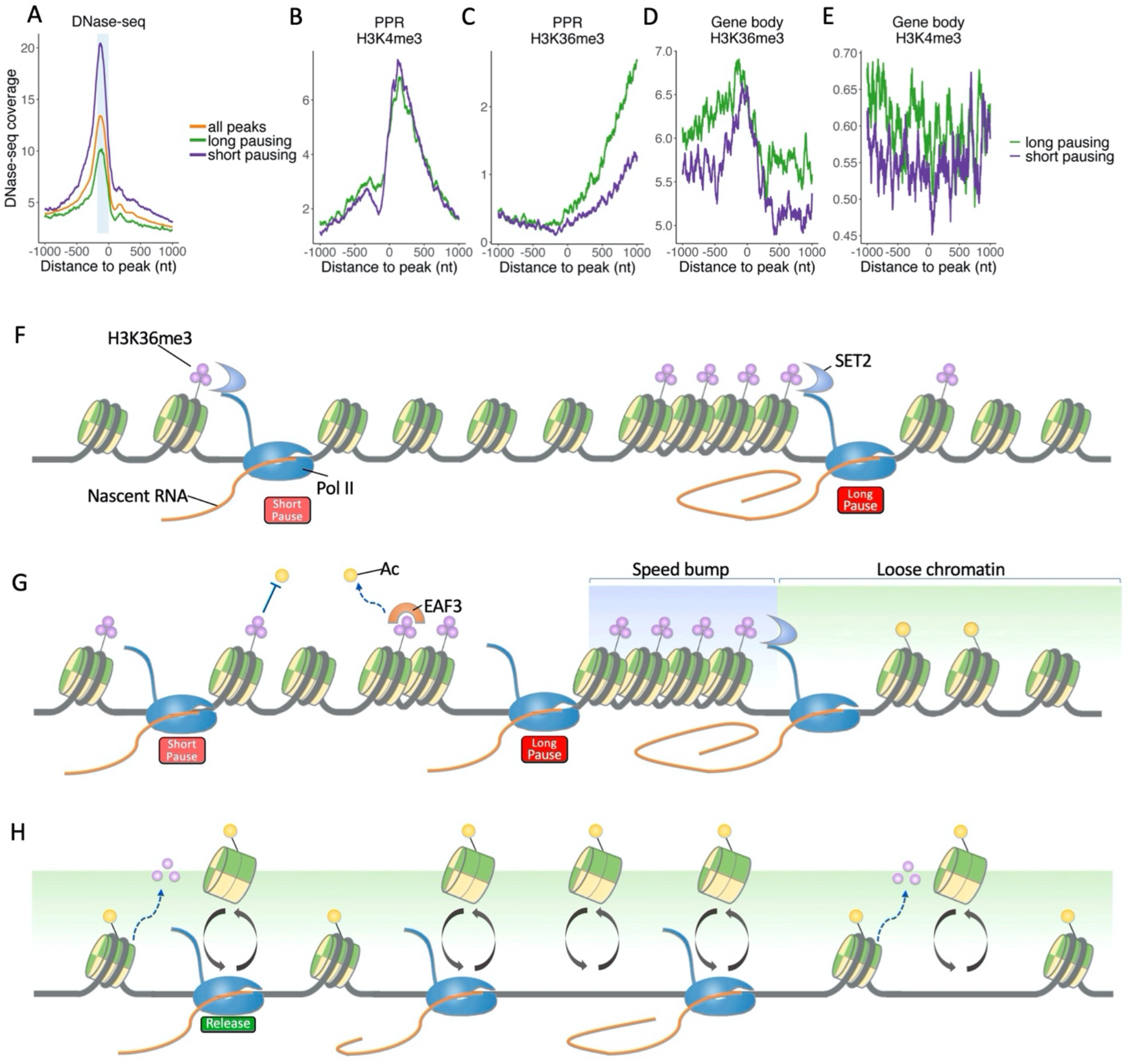
Chromatin state and pausing times. (**A**) Peaks were classified into ‘long’ and ‘short’ according to their pausing times. The average signal of DNase-seq data is displayed in the vicinity of the two classes of peaks (and all peaks). The region from −180 to the peak was shaded in light blue. (**B**) Peaks were classified similarly as in (A); signal profiles of H3K4me3 ChIP-seq data of peaks within first 500bp of gene are shown. (**C**) Similar to (B), for H3K36me3. (**D**) Similar to (B), for the peaks within the gene body (Except the first 2000bp and last 1500bp of gene). (**E**) Similar to (D), for H3K4me3. (**F**) CTD of paused Pol II recruits SET2 to methylate H3K36. H3K36me3 level increases if the pausing lasts longer. (**G**) Model of the dynamic equilibrium between H3K36me3 and histone acetylation under homeostasis. Two H3K36me3 related pausing sites have been set. Packaged H3K36me3 can form into a ‘speed bump’ which establishes long pausing, while shorter pausing might correspond to isolated marks. The Pol II CTD can recruit SET2 to methylate H3K36. The H3K36me3 then can facilitate deacetylation of histones (by active EAF3) and/or inhibit histone acetylation. (**H**) Histone acetylation releases paused polymerase after removal of H3K36me3, resulting in a transcriptional burst.

Activating histone modifications (58) such as H3K4 methylations and H3K27 acetylation exhibit similar profiles around long and short pausing sites within the PPR (Fig. 6B and Fig. S10). H3K36me3 is an elongation marker that is usually found enriched at exons of active genes (63, 64). In contrast to other active markers, H3K36me3 shows a clear pattern of enrichment downstream of long pausing sites in the PPR (Fig. 6C) and is also enriched at the pausing sites in the gene body (Fig. 6D). This suggests that H3K36me3 is involved in pausing, specifically, long pausing. This contrasts with other activating histone modifications, whose profiles appear flat in gene bodies despite having higher coverages around long pausing sites (Fig. 6E and Fig. S10). Its association with active genes could mean that H3K36me3 is involved in the more intensive pausing activity we find in highly expressed genes (Fig. 4A-B).

Two hypotheses could be posed for explaining the reason that H3K36me3 is an active marker of expression but also associates with long pausing. The first one posits that pausing could help the recruitment and function of the SET2 complex. Since H3K36me3 is deposited co-transcriptionally, increasing Pol II residence time would also give SET2 more time to act. The longer the pausing lasts, the higher the methylation level of H3K36 would thus be expected to be (Fig. 6F). The second explanation is that the mark is deposited in the wake of elongating Pol II rather than functioning as a pre-set, static marker. Methylation of H3K36 is carried out co-transcriptionally by the SET2 complex which is recruited by the carboxy-terminal domain (CTD) of Pol II (65). By facilitating histone deacetylation via activation of EAF3 (66) and remodelling of repressive chromatin (67, 68), H3K36me3 might thus act as a ‘speed bump’ to prevent collisions between succeeding polymerases (Fig. 6G). This would also explain why a loss of SET2 only slightly influences expression levels of H3K36me3 positive genes (69). Interestingly, a longer continuous H3K36me3 region in the gene body will form into a stronger speed bump which blocks polymerase for a longer time (Fig. 6E). A tug of war between H3K36me3 and histone acetylation may function as speed control for elongation: paused polymerase is released by demethylation of H3K36me3 and histone acetylation in response to stimuli such as heat shocks (44), thus raising the elongation rate of polymerase (Fig. 6H). These two mechanisms could also function together.

In order to further gauge the quality and informative value of these TV-PRO-seq based results, we carried out a side-by-side comparison with NET-seq data for the same cells and chromatin states in different regions. NET-seq shows completely different patterns for H3K36me3 in the PPR compared to TV-PRO-seq (Fig. S11). The reason is probably that its signal not only correlates with pausing time, but also with pausing fraction, abortive transcription and expression level (Fig. 2A-D, Box1). As H3K36me3 has been identified as elongation marker, high polymerase occupancies are expected for genes with high H3K36me3 loads. The high polymerase occupancy contributes to a low PI and leads to opposite results for TV-PRO-seq and NET-seq, further demonstrating our assay’s benefit.

## Discussion

While the phenomenon of polymerase pausing has been known for decades (3, 16), the methods to investigate it are still mainly based on polymerase occupancy (3, 8, 24, 29), which can be actually confounded by various other factors.

Highly expressed genes tend to have more polymerases on their bodies regardless of the pausing, therefore a non-pausing position of a highly expressed gene can have an even higher polymerase occupancy than a pausing site of a low-expression gene. As all the positions in a gene are associated with the same expression level, using the polymerase occupancy of non-pausing sites to normalize the occupancy of pausing sites can reduce the influence of the expression level towards the occupancy.

It has been suggested that the high density of polymerases in PPRs is due to long pausing in these regions (6). However, this suggestion is based on the assumption that all nascent transcripts from a gene are eventually expressed as full length RNA and/or share TSS and TES. In fact, genes have wide transcription initiation domains (26) and most transcription terminates before entering productive elongation (20–22). This means that only a fraction of polymerases will reach a gene’s 3’ end, leading to the high polymerase occupancy observed in the PPR. Based on our findings, we propose that the activity of pausing sites can be regulated. When the pausing site has been turned off, polymerase can pass it unimpededly. A pausing site with high pausing fraction and low pausing time can thus have the same average residence time as a pausing site with low pausing fraction and high pausing time (Box 1, Fig. 2A-D). Since TV-PRO-seq compares reads from the same positions, it can remove the influence of the polymerase flux. Furthermore, since non-pausing polymerase makes only tiny contributions to the polymerase occupancy (Fig. 2D; Box 1), TV-PRO-seq can also reduce the influence of the pausing fraction and measure pausing time of paused polymerase only. Our results suggest that NELF can stabilize pausing (Fig. S6, 3E) and that polymerase indeed frequently pauses in the PPR. However, the median pausing time of individual pausing sites in PPR is less than the other regions’ peaks’ (Fig. 2E).

Unlike measurements from Trp-related methods, which are largely limited to study the PPR, TV-PRO-seq yields results at single nucleotide resolution, genome-wide (Fig. 1A-B). This advantage allowed us to analyse pausing times in much greater detail and bigger scale than what was previously achieved. We found that, unlike peaks in the first 120bp of genes, pausing in the +180 to +320 region appears resistant to sarkosyl treatment (Fig. 3A). Also, we were able to investigate pausing times at locations far from TSSs; TV-PRO-seq revealed pausing times of peaks further downstream within genes and even beyond the genes’ 3’ ends (Fig. 2F). Trp-based studies would not be able to easily show this due to their focus on the PPR (Fig. 1B).

Beyond these Pol II related analyses, TV-PRO-seq is suited to examine transcription by other RNA polymerases. Exploiting this potential revealed that Pol III has significantly shorter pausing times than Pol II and Pol I (Fig. 1F). This highlights the relevance of pausing for transcription by the two former polymerase types, too; upon closer inspection of tRNA genes, a distinct pausing pattern emerged indeed (Fig. S5A).

TV-PRO-seq also allowed us to integrate the pausing time profiles with other genome-wide data. By grouping genes with different expression levels and different transcriptional noise levels according to droplet single cell RNA-seq data (53), we were able to uncover an intriguing relationship between expression and pausing, that is, highly expressed genes not only have more pausing sites (Fig. 4A), but also longer pausing times at each of these (Fig. 4C). For noisier genes, we find both more (Fig. 5A) and longer pausing (Fig. 5C); this is consistent with predictions of modelling works (52, 56, 57). We further integrated pausing times with ChIP-seq data of histone markers and found the active transcription marker H3K36me3 to correlate with pausing in the gene body (Fig. 6E). We propose that paused polymerase could recruit SET2 for methylation of H3K36 and/or H3K36me3 pauses polymerase by repressing histone acetylation (Fig. 6F-H).

In summary, TV-PRO-seq provides a powerful novel tool to time polymerase pausing. It permits genome-wide estimation of pausing release times at single base resolution. Our analyses illustrate the rich new insights that can be obtained with our approach in regard to the different polymerase types, the dynamics associated with the different pausing sites, the chromatin state, and, more generically, the process of stochastic transcription. These findings would be hard to obtain with competing techniques, such as NET-seq, which reflect only the polymerase occupancy. Our data provide promising starting points for further investigations, including the study of pausing at short genes such as tRNA and lncRNA loci, the mechanisms involved in pausing, and several other related subjects.

## Limitations

TV-PRO-seq is based on PRO-seq; it can only reflect pausing profiles in vitro as the nucleotide run-on of biotin-NTP is performed in permeabilizated cells. Plus, polymerase that is paused in the region very close to TSSs cannot be fully detected as short reads are removed before alignment.

TV-PRO-seq requires preparation of ≥4 different parallel PRO-seq samples with independent cell permeabilization and library building efforts; this is a source of noise in terms of sample variation, along with technical noise that commonly affects PRO-seq data quality. As TV-PRO-seq provides single base resolution on genome-wide scale, even high sequencing depths will result in relatively low read counts on individual pausing sites, limiting sensitivity and precision. This effect can be alleviated, though, by considering and analysing ensembles of pausing sites.

TV-PRO-seq disallows sarkosyl (which artificially facilitates polymerase release via the run-on buffer) thus making library building more demanding than conventional PRO-seq.

Three hypothetical pausing sites have been set for illustration. The grey line represents a template DNA which includes three examples for pausing sites. The red points denote pausing events that occurred during transcription.

NET-seq, PRO-seq and ChIP-seq reads reflect polymerase occupancy in a broader sense. The polymerase occupancy is influenced by various factors.

The **polymerase flux** is the number of polymerases that move past a given position in a unit of time. The polymerase fluxes in different regions of a gene are not necessarily equal; for instance, abortive transcription increase the polymerase flux in the PPR compared to downstream. Polymerase flux is affected by backtracking and spurious transcription and impacts the gene’s expression level, but cannot be easily inferred from it.

The **polymerase occupancy** (which is the number of polymerases at a given genomic position) equals the polymerase flux times the average residence time of polymerases:

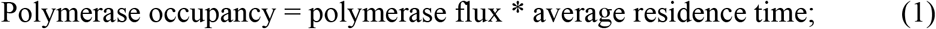

Not all polymerases will stop when they encounter a pausing site. We refer to the average fraction of polymerases that actually pause at a pausing site as the **pausing fraction**.

If a polymerase is elongating, the residence time of polymerase at each position is very short and contributes a small amount to average residence time measurements. If we neglect it, we obtain:

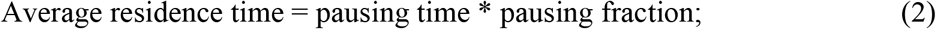

The **pausing time, which is the residence time of paused polymerase,** can be measured by TV-PRO-seq.

Combining (1) and (2), we get:

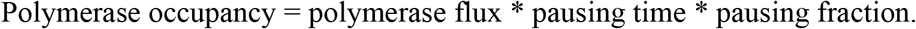

Finally, we refer to the **pausing frequency** as the number of pausing sites per unit of length.

## Materials and Methods

### Time-variant PRO-seq library building

HEK293 cells were grown to 60% confluency at 37°C and 5% CO_2_ in a 175 cm^2^ flask in DMEM supplemented with 10% FBS. One day before permeabilization of cells, the culture medium was replaced with fresh medium. For Trp treatment, Trp was added at a concentrations of 500 nM, then cells were incubated at 37°C for 10 min, followed by permeabilization.

Cell permeabilization was carried out following the PRO-seq protocol (30). Permeabilized cells were stored at −80°C. The cells were placed on 37°C for 3 min for thawing. Thawed cells were further processed by adding biotin-labeled NTPs. Two replicates of 4-biotin run-on samples were prepared for HEK293 cells following the PRO-seq protocol. Furthermore, duplicates of 4-biotin run-on samples were prepared in HEK293 cells in a run-on buffer without sarkosyl. The main TV-PRO-seq experiment consisted of 4 independent PRO-seq samples of the 4 run-on times 30 sec, 2 min, 8 min and 32 min. After run-on, the experiment followed the PRO-seq protocol (30). For the Trp treatment sample, 8 min run-on with sarkosyl was performed.

### Processing of sequencing data

Sequencing was performed on an Illumina NextSeq 500 for 51bp single end. Raw data was converted into FASTQ format by bcl2fastq with 0 index mismatches allowed.

Reads were trimmed with Cutadapt version 1.14 (70), to remove sequences starting with the adaptor sequence ‘TGGAATTCTCGGGTGCCAAGG’ from the 3’ end of reads, and reads shorter than 20bp after trimming were discarded:

~~~
  cutadapt -a TGGAATTCTCGGGTGCCAAGG -m 20 -e 0.05
~~~

Trimmed reads were aligned to the best matched position of hg38 genome with Hisat2 version 2.1.0 (71), resulting in alignment rates above 80%:

~~~
  hisat2 -p 4 -k 1 --no-unal -x ~/hg38/genome -U data_2.fastq.gz -S data.sam
~~~

Because the ends of sequencing reads have lower sequencing quality, Hisat2 uses soft clipping for the reads, which moves the detected pausing site upstream of the actual pausing site. A custom script `Sam_enlong.pl` was used on the SAM files to extend the soft clipped reads to their original lengths.

Because sequencing depth also has an influence during the process of peak calling of TV-PRO-seq, another script `Sam_cutter.pl` was used to reduce the 4 TV-PRO-seq SAM files for each PRO-seq sample to the same sizes by randomly selecting a subset of reads for each.

The processed SAM files were further converted to BAM files and were sorted with samtools version 0.1.19 using `samtools view -S -b and samtools sort` (72).

The sorted BAM files were then converted to BEDGRAPH files (73). The 5’ end of a read corresponds to the position of the paused polymerase release site on the opposite strand:

Pausing on plus strand: `genomeCoverageBed -strand - -5 -bga -ibam`
Pausing on minus strand: `genomeCoverageBed -strand + -5 -bga -ibam`

We then combined the BEDGRAPH files for the various replicates and time points into two files, one for each strand, with the custom script `TV_bedGraph_merger.pl`. These files corresponded to tables with rows for each position and columns containing the read numbers across the samples, and were used for the further analysis.

### Peak calling

We developed a custom procedure for peak calling from single-base resolution strand-specific sequencing experiments such as TV-PRO-seq. Rather generically, we require that the transcription level *μ* at a peak exceeds a threshold value *Q_bio_* which depends on local fluctuations:

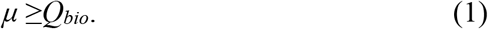

The actual procedure is based on the aggregated reads from all the experiments at different run-on times and for a specific position (hereafter, such total reads per nt will be simply referred to as the “total reads”) and is detailed below.

1. A threshold *t* for the minimum number of reads on each single genomic position was set. More precisely, genomic positions with total read higher than *t* were selected as ‘candidate peaks’ for further analysis. The basic threshold *t* has been heuristically set to 10 and will vary with sequencing depth. In addition to this, we discard the candidate peaks if the number of reads is zero for all the replicates corresponding to a single one run-on time, at least.
2. Secondly, we address the fact that some polymerase pausing regions are wider than one nt (8). An example of such a dispersed pausing region is illustrated in Figure S1, within a 50-nt fragment of plus strand of chromosome 1. In Figure S1, we consider the position with most reads in the dispersed pausing region. To deal with this, we exclude a ‘candidate peak’ if another ‘candidate peak’ has more reads in its +/− three-nt neighborhood. This ensures that only a single position is selected from a dispersed peak.

For highly expressed genomic regions, it is likely that some positions have a large number of reads (viz., higher than the threshold *t*) and pass selection step 1, even if they correspond to regions with constant elongation rate and do not have significant pausing. Similarly, along the same non-pausing regions, the step 2 returns the genomic positions that have the highest amount of reads, even if this is just due to random fluctuations. As an example, the genomic positions marked as purple in the fragment illustrated in Figure S1 corresponds to such a case. Therefore, a third step is necessary to filter the candidate peaks that are likely to be located in a region of constant elongation rate but cannot be discarded during the steps 1 and 2. We perform a two-step procedure as explained below.

3.1. The first sub-step consists of assessing the local biological fluctuations in the polymerase occupation and deriving the threshold *Q* of condition (1). We assume that the polymerase occupancy in a constant elongation-rate region follows the Poisson distribution with parameter *b*. As the average elongation rate across the mammalian genome is about 33.3 nt/sec (28), we expect that, in such non-pausing regions, all the polymerases are released by the time of the first run-on experiment (i.e., 30 seconds); therefore, for these regions, the differences observed between experiments at different run-on times are presumably due to statistical fluctuations, suggesting that we can actually ignore the dependence on run-on time and aggregate the reads across all experiments. We then focus on the reads across the +/−100-nt neighborhood around each candidate peak. Their mean reads, averaged over both the replicates and the 201 nts, yields the expected number of reads *b* per nt^1^ (in the neighborhood). Based on a null local Poissonian assumption, as if reads were Poisson distributed with rate *b*, we associate an upper *qth* quantile *Q_bio_* to each neighborhood, where the value of *q* is heuristically chosen to control the number of (false positives) bases whose read number exceeds *Q_bio_* purely due to statistical fluctuations. Our (rather conservative) choice would be to allow only one false positive in the whole ‘active genome’. We define the latter as all positions with at least one read. Since from our experiment there are 111868728 such bases, we heuristically set *q*=1/111868728.
3.2. Secondly, we need to assess the sequencing noise as a function of the transcription level. To this end, we sequenced one of the replicates (specifically, the second 32-minute run-on replicate) twice, and trimmed the technical replicate with the highest total alignment reads to the same level as the other one. This trick gave us two replicates of identical total aligned reads, from which we computed the average reads for each nt. Further, by gathering the positions whose average read equals a certain number *μ* and computing their CV^2^ we obtain the scatter plot of Figure S1B, which appears to closely follow the fitted standard noise model CV^2^ = *A/μ + B*, and which can be expressed as

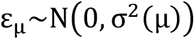

where

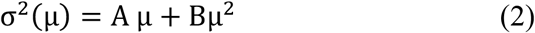 As an example, see Figure S1B for the empirical distribution of the reads centered at *μ*=20 alongside its Poisson and normal fit). Based on this model, the (observed) peak read is randomly drawn from

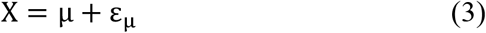

from which it follows that selecting the candidate peaks with more reads than the 0.99th quantile *Q_seq_* of the normal distribution centred at *Q_bio_* with variance *σ^2^*(*μ*) satisfies condition (1) with probability 0.99,

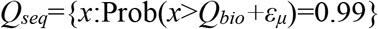 Since we don’t know the value of *μ* to insert into equation (2), we replace it with either *Q_bio_* or the peak reads itself; the first choice underestimates *Q_seq as_ Q_bio_* < *μ* (for all the non-trivial cases) and hence σ^2^(*Q_bio_*) < σ^2^(*μ*), while the second choice has not such a bias as *X* is centred at *μ*. It is worth noting that there is an alternative but equivalent choice: one can compute the lower quantile of the distribution centred at the peak read *x, Q’_seq_*={*q*: Prob(*q* < *x*+*ε*)}, and require that *Q’_seq_* > *Q_bio_*. In conclusion, we incorporate the polymerase noise model of point 3.1 and the sequencing noise model of point 3.2 into condition (1) by choosing the candidate peaks such that *x* ≥ *Q_seq_*, where *Q_seq_* depends on *Q_bio_*.

### Inference of single-nucleotide transcription rates

In this section, we derive a simple Bayesian model for TV-PRO-seq data and detail the procedure to infer the single-nucleotide transcription rates β_i_. We are interested in the stochastic dynamics of biotin-NTP incorporation into a nascent mRNA which can be represented as the following simple reaction:

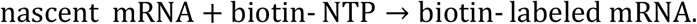

Such a reaction corresponds to one transcription step and is specific to the genomic position i complementary to the 3’-end nucleotide of the nascent mRNA. Assuming that the biotin-NTP population is large and remains constant during the reaction progress, we obtain

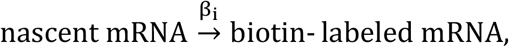

which occurs at constant single-nucleotide transcription rate β_i_. The average time that the PolII spends on the base i is the reciprocal 1/β_i_, which we refer to as the *pausing time*.

Let y_i_(t) and x_i_(t) denote the average populations of nascent-mRNA and biotin-labelled mRNA (specific to the genomic position i), respectively. The following rate equation is satisfied:

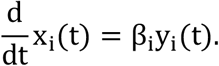

As the presence of the biotin prevents further elongation and no new transcription is initiated, y_i_(t) naturally decays according to

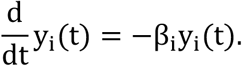

Solving this simple system of ODEs with initial conditions

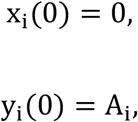

yields

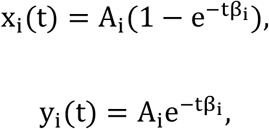

which predicts that the average population of the biotin-labelled mRNA increases up to the saturation point A_i_ while the unlabelled nascent mRNA is depleted according to exponential law.

Our analysis focuses on a subset of genomic positions i ∈ S, which we refer to as *peak* positions, where transcription level saturates to A_i_ at rate β_i_. We speculate that a large number of genomic positions displays negligible pausing with Pol IIs stepping forwards shortly after biotin-NTP treatment and with transcription level concentrating around A_bck_. We refer to such positions as *background*. Therefore, the expression level of the whole genome x_tot_(t) = ∑_i∈s_ x_i_ (t) + x_bck_(t) grows according to

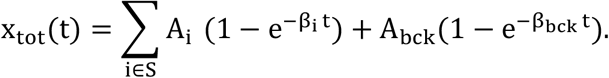

While we have a model for the average transcription level x_i_(t) at genomic position i ∈ S and run-on time t, the average number of reads N_i_(t) depends on the sequencing depth κ(t) which is different for each sequencing experiment and therefore depends on the run-on time t, i.e.,

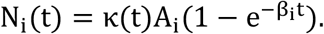

It is convenient to study the ratio x_i_ = N_i_(t)/N_tot_(t), where N_tot_(t) = κ(t)x_tot_(t), as the dependence on κ(t) cancels out. This represents the expected number of reads from the region of interest (e.g., from a peak position) normalised by the average total-genome reads at the same run-on time t.

We obtain the normalised model

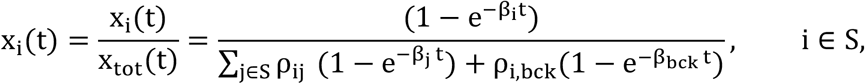

where ρ_i_ = A_j_/A_i_ and ρ_i,bck_ = A_bck_/A_i_. We will later consider an approximated model where the growth curve X_tot_(t) is described by a single effective rate β_tot_.

The quantities x_i_(t), i ∈ S, can be organised into an |S| × T matrix X where T is the number of predictor observation run-on times. This allows us to use the compact notation

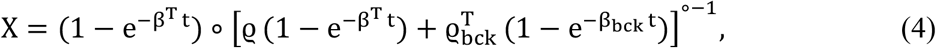

where t = (t_2_, t_2_, …, t_T_) is the vector of predictor observation run-on times, β = (β_1_, β_2_,…, β_|s|_) is the vector of rates, 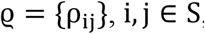, and 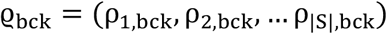 incorporates the relative saturation points. The notation A ◦ B is the Hadamard (element-wise) product of A and B while A^◦−1^ is the Hadamard inverse of A.

To simplify this model, we use the naïve form

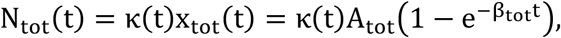

which approximates the growth of the average of total reads. The total number of mitochondrial reads xchrM appears to saturate much quicker than the pausing site reads and, to a first approximation, we assume that *x*_chrM_ = *κ*(*t*)*A*_chrM_. We divide the total reads by the chromosome-M reads, and fit the model

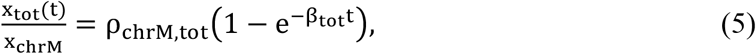

where ρ_chrM,tot_ = A_tot_/A_chrM_, to such data using the random-search algorithm of the nls2 R package (74), which returned a fit with estimated parameters reported in the table below.

|  | Estimate | Std.err. | t value | Pr(>\|t\|) |
| --- | --- | --- | --- | --- |
| $\rho_{\text{chrM,tot}}$ | 14.993 | 1.230 | 12.192 | 0.000 |
| $\beta_{\text{tot}}$ | 4.837 | 1.985 | 2.437 | 0.050 |

Our choice is to use the exponential model to approximate the growth of the average total-genome reads N_tot_(t), and study

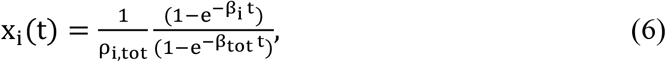

where i ∈ S and ρ_i,tot_ are parameters fixed by data. In matrix form, we get

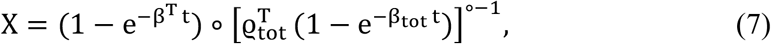

where

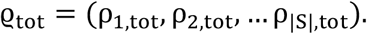

Dividing equation (6) by (1 − e^−*β*_tot_*t*^) yields the more intuitive saturation curves of Figure 1A. We then chose the informative prior

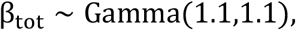

where Gamma(α, β) represents the Gamma distribution with mean α/β and variance α/β^2^, which places substantial mass around 1 and little mass around 0^+^. The peaks must have an average rate of the same order as the total growth rate, although the growth rates corresponding to pausing elements can be significantly smaller. Based on these considerations we chose the informative priors

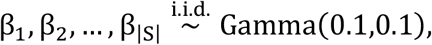

which have mean and variance equal to 1 and 10, respectively, and place lot of mass at 0^+^.

The next steps consist of incorporating noise and thus defining a Bayesian model to be fitted. We incorporate the noise in the model as follows. The sequencing reads are obtained after several amplification steps and are restricted to be positive. Hence we assume that the observables Y are subjected to multiplicative errors with lognormal distribution, i.e.,

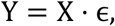

where

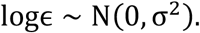

As ϵ = e^σZ^ with Z ~ N(0,1), we get

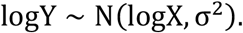

To empirically guess a prior distribution for σ given the coefficient of variation of Y, we use the error-propagation formula

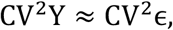

where CV^2^Y is estimated from aggregated data. As ϵ is lognormal, we have

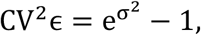

and

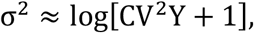

which suggests the prior

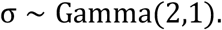

An MCMC sampler to fit the model was implemented using the PyMC3 Library for Bayesian Statistical Modeling and Probabilistic Machine Learning (75). PyMC3 relies on the Theano framework (76), which allows fast evaluation of matrix expressions, such as those in equations (4) and (7), and offers the powerful NUTS sampling algorithm to fit models with thousands of parameters. Nevertheless, we aim to infer the growth rate of up to ~ 60000 peaks. To ease the computational burden, we divide the peak list into chunks of ~ 3000 randomly chosen peaks. The simulations were performed on CyVerse computational facilities (77). Further, we averaged the reads over the replicates, and the averages at 32 minutes of run-on time are used as saturation levels.

In addition to the estimates of the peak rates, the method returns estimates of β_tot_ from each chunk. These are very close to the rate 0.1min^-1^ obtained from the half-life measured in Jonkers, Kwak, and Lis (28). Aggregating the individual-chunk estimates using the laws of total mean and variance yields:

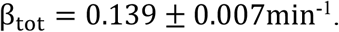

In order to assess the sensitivity with respect to the prior distribution, we also ran the inference procedure using the vague prior distributions:

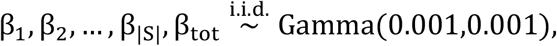

which results in a wider range of inferred β_i_, whilst conserving the overall rank order.

### Peak annotation to 3’ and 5’ ends of exons

Two reference lists were used for annotating the ends of the target regions. For mRNA genes, the list was downloaded from UCSC table browser with parameters: assembly - hg38, group - mRNA and EST, table - UCSC RefSeq, output format – All field from selected table (78). The 5’ and 3’ends of all exons from the mRNA list was transformed into another table with the custom script `Unique_annotation_maker.pl`. Column 1 to 3 was the chromosome, position and strand of annotation site; column 4 was the gene name; if the column 5 ‘type’ equals ‘start’, it means it is the 5’ end of exon, otherwise it is 3’ end; the column 6 ‘number_min’ and 7 ‘number_max’ are the min and max number of exons in different variants of the same gene, respectively, and the TES are marked as −1; The column 8 ‘hit’ shows how many variants of a transcript have this splicing site and column 9 ‘variant’ refers to the number of transcript variants the gene has.

For rRNA and tRNA genes, two tables were downloaded from RNAcentral (*https://rnacentral.org/*), the RNA gene classification information was in rfam_annotations.tsv and the genomic locations of these genes was in Homo_sapiens.GRCh38.bed, respectively. We used a custom script `rFAM_annotation_merger.pl` to merge these two tables for the further analysis (79).

The two annotation files were used for annotating peaks by another custom script `Peak_annotater.pl`, which identifies peaks located within a specified distance of the annotation site. For example, we can detect the peaks located in a ±4500 nt region of all the 5’ and 3’ ends of UCSC refgene mRNA genes with the following command:

~~~
  perl Peak_annotater.pl All_mRNA Beta_summary 4500
~~~

The peaks that were annotated to have ‘type’ equal to ‘start’, ‘number_max’ equal to 1 and ‘hit’ equal to ‘variant’ were those near the TSS of genes with unique TSSs. The sense and antisense reads around these unique TSSs were used to generate the density plot using the ggplot2 package (80) for *R* (Figure 1B).

### Peak annotation within genic regions

For mRNA transcripts by Pol II, `UCSC2bed.pl` was used on the same UCSC list as above, and for rRNA transcripts by Pol I, the script `rFAM_region.pl` was used for transforming the merged list from RNAcentral. Pol III target regions were taken from published data (39); we used the ‘Potential Pol3 targets’ table and converted it to human genome assembly GRCh38 with the UCSC liftOver tool (78). The output BED file contained 6 columns: chromosome, start of region, end of region, gene name, gene type/transcript ID and DNA strand.

The custom script `Annotation_region.pl` was used to extract peaks in the target regions according to the annotation lists generated above. The peaks annotated by Pol I, Pol II and Pol III were compared to the peaks detected on chrM in terms of their pausing time distributions. These were displayed as violin plots with inserted boxplots using the ggplot2 package for *R* (Figure 1G).

### Metagene analysis about pausing peaks

15993 genes which have unique TSSs and TESs and are longer than 3000 nt were used for metagene analysis. We classified the peaks into 7 regions: 1. Promoter, 2. TSS related region, 3. earlier intron, 4. exon, 5. later intron, 6. region before TES and 7. pA related region.

We obtained regions 1, 2, 6 and 7 from the annotations of 3’ and 5’ ends of exons from the list generated with `Peak_annotater.pl`.

Promoter: 1000-nt region upstream of TSS
TSS related region: 1000-nt region downstream of TSS
region before TES: 500-nt region upstream of TES
pA related region: 4500-nt region downstream of TES

The peaks in the introns and exons were annotated with `whole_gene_annotater.pl`, using the annotation list generated with `whole_gene_annotation_list_maker.pl`. Only exons and introns not overlapping with the first 1000-nt or last 500-nt of transcripts were selected. If the intron’s centre position was in the first half of the gene, we considered an intron to be an early intron. Otherwise we regarded it as a later intron.

Because most exons or introns have different lengths, we normalized the peak densities before plotting. First, the peaks in introns and exons were annotated with the relative location, that is the distance between the peak and the 5’ end of the annotated region, divided by the length of the annotated region. Then we calculated the average length for each region and multiplied it with the relative location.

To show the pausing times of the 7 regions defined above, a smoothed conditional mean plot with LOESS fitting was generated using the ggplot2 *R* package with parameter span=0.1 for the ggplot function (Figure 2E, F). We also separately plotted the smoothed conditional mean plot for peaks around TSSs (Figure 3A). Peaks around TSS and TES of tRNA genes were plotted in the same way (Figure S5).

### Gene expression level and transcriptional noise estimation and selection

Genes’ expression levels were calculated as average UMI counts from single-cell sequencing data (54). Genes within the top 20% of expression levels were identified as highly expressed and the bottom 20% as less expressed.

To estimate transcriptional noise, we computed the ‘above Poisson score’*η* of genes from single-cell sequencing data as in (54):

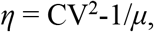

where *μ* is the mean mRNA number for a gene, and CV is its coefficient of variation. We selected genes with the highest and the lowest noise heuristically, taking into account the dependence of *η* on *μ* as follows. We processed the single-cell sequencing dataset of (53) with the custom script `Rank_eta.pl`. This first sorts the genes into a list by their mean expression. It then moves a sliding window of size *WS* = 100 along this list and, at each position of the window, ranks the genes with regards to the value of *η* and records these ranks. For each gene in the list, a number *WS* of ranks results, of which the top and bottom ranks are averaged to give the ‘noise score’. We refer to genes within the top and bottom 5% noise scores as ‘high noise’ and ‘low noise’ genes. For genes with equal noise scores, this procedure was repeated for *WS* = 20 and *WS* = 500, and rescaling the resulting noise scores to the range 0 to 100, followed by averaging across the three noise scores (Figure S8).

We generated the smoothed conditional mean plots of the ‘highly expressed’ and the ‘less expressed’ genes of Figure 4D and the ‘high noise’ and ‘low noise’ genes of Figure 5D using the same strategy as in the full metagene analysis (Figure 4D, 5B) and plotted histograms to show the absolute frequencies of peaks from ‘high noise’ and ‘low noise’ genes (Figure 4A, 5A). Split violin plots (Figure 4C, 5D) were generated with ggplot2 as before.

### Histone modification and chromatin accessibility for TV-PRO-seq data

We used existing HEK293 cell ChIP-seq data for different histone modifications from published studies and/or public depositories for the analysis. NELF data were obtained from Gene Expression Omnibus, GSE109652 (81); H3K4me1, H3K4me2, H3K4me3 and H3K27ac data were obtained from GSE101646 (82); H3K36me3 and DNase-seq data were downloaded from ENCODE series ENCSR372WXC and ENCSR000EJR. The data were first trimmed with Trimmomatic-0.36 with options `LEADING:24 TRAILING:24 SLIDINGWINDOW:4:20 MINLEN:20` (83), then aligned to hg38 under --no-spliced-alignment condition by Hisat2 (71). The SAM files were converted to BAM files, then to BED files using Samtools (72) and Bedtools (73), respectively. The read intervals in the BED files were adjusted to the same lengths with the custom script `bed_normal_length.pl` to make sure the coverages of reads bore equal weights for each read. We then converted the data to BEDGRAPH files with the `genomeCoverageBed` command from Bedtools, using the flags `-bga` (73). The BEDGRAPH files were annotated to TSS or pausing peaks with the custom script `Liner_bedgraph.pl`.

We then classified peaks on nuclear chromosomes into those with the longest 5% and shortest 5% pausing times, and extracted the coverage from the BEDGRAPH files within +/- 1000 nt of each peak in both classes. We then removed the top 5% of these coverage intervals since these had disproportionately strong influence on the results. The peaks were further classified by the position within genes. PPR refers to the first 500bp of a gene and the gene body to the region after +1500 from TSS and before −1500 from TES. Finally, we averaged the coverages of each class, respectively, and displayed the results using ggplot2 in *R* (Figure 6A-E, Figure S10).

The NELF levels of peaks were defined as the average ChIP-seq coverage in the region +/- 80bp of peaks. The peaks within the top 10% of NELF levels were identified as high NELF level and the bottom 10% as low NELF level.

### Histone modification and chromatin accessibility for mNET-seq data

HEK293 mNET-seq data were downloaded from Gene Expression Omnibus, GSE61332 (84). We used the UCSC liftOver tool to convert the BEDGRAPH file to hg38 (78). We then defined target genes for further analysis by selecting genes longer than 3000 nt, with unique TSSs and TESs. Peak selection for the mNET-seq data followed the same strategy as for TV-PRO-seq; The peak selection output file was processed with the script `Liner_bedgraph.pl` to extract histone modification states within +/-1000 nt of peaks in the same way as for TV-PRO-seq. We estimate pausing of mNET-seq by the PI (pausing index), which divides the read numbers of peaks by the average read numbers in the gene body (+500 to TES). We removed the top 5% peaks with the highest average coverage of each group and plotted the average coverage of histone modification at peaks corresponding to the top and bottom 5% PI, respectively (for all peaks in target genes, peaks within the TSS to +500 region only, or peaks within the +1500 to TES region only).

In order to compare TV-PRO-seq and mNET-seq with regards to the chromatin state results, we needed to subset the TV-PRO-seq data to the same target genes as we used for the mNET-seq data. The script `PI_TV_annotater.pl` was used to extract the coverage information of individual TV-PRO-seq peaks located in the target genes. We then selected long pausing and short pausing peaks as above. The average ChIP-seq/DNase-seq coverages of long pausing and short pausing peaks were then used for comparison with the high PI and low PI peaks (Figure S11).

## Acknowledgments

**General**: We thank Andrew Nelson and Keith Leppard for reading the manuscript and making valuable suggestions. This work was supported by BBSRC grants BB/L006340/1 and BB/M017982/1, and EPSRC grant EP/T002794/1. Parts of the work were carried out by MC during an earlier affiliation with the Department of Statistics, University of Warwick.

## Author contributions

JZ designed the study and carried out experimental work. JZ, MC and DH analysed the data, carried out theoretical work, and wrote the manuscript. DH supervised the work.

## Data and materials availability

All data needed to evaluate the conclusions in the paper are present in the paper and/or the Supplementary Materials. Sequencing data generated in this study was deposited in GEO under the accession number GSE118957.

## Supplementary Materials

**Fig. S1.**
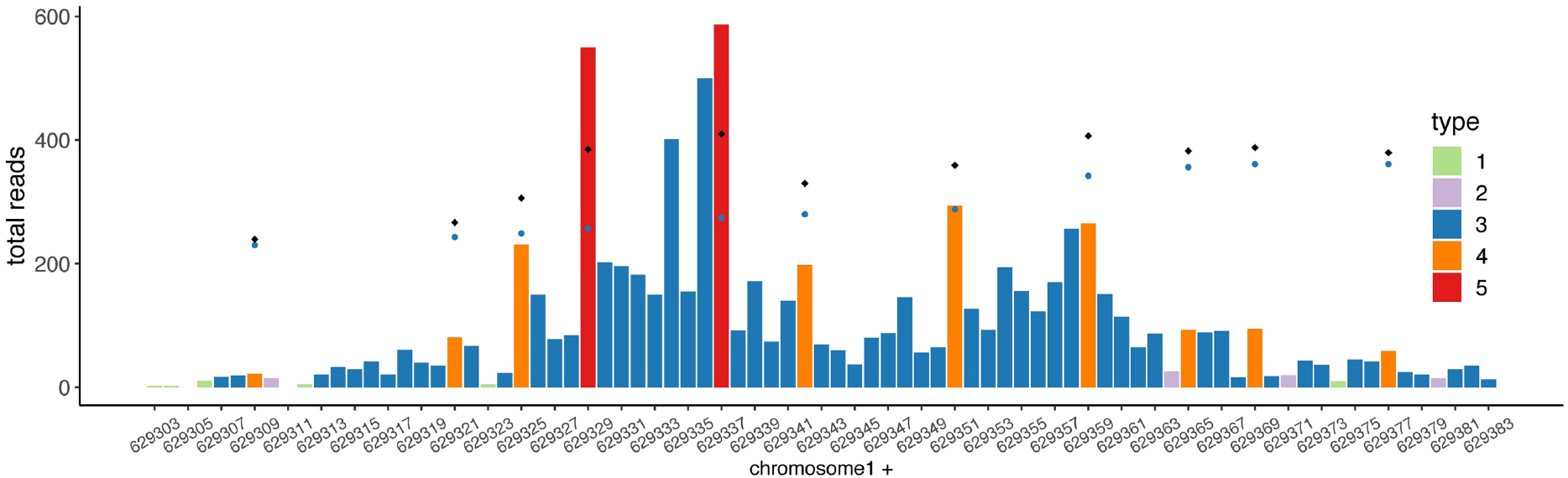
Peak selection. The total reads from an 80-nt fragment of the + strand of chromosome 1 are shown as an example of the peak selection procedure. Positions of color 1 (green) are discarded at the first step as they have less than 10 total reads, and positions of color 2 (purple) are rejected for having zero reads in at least a single run-on time sample. Positions 629336-6329337 correspond to a 2-nt dispersed peak, where only the first base is selected for further analysis, while position 629329 corresponds to a peak concentrated on a single base. Peaks of color 3 (blue) are discarded at step 2 in favour of the highest peak in the +/-3-nt neighborhood. Peaks of color 4 (orange) are discarded at step 3. Red peaks (color 5) are selected/called for further analysis. Blue points correspond to the quantile threshold Q_bio_. Black points correspond to the threshold Q_seq_.

**Fig. S2.**
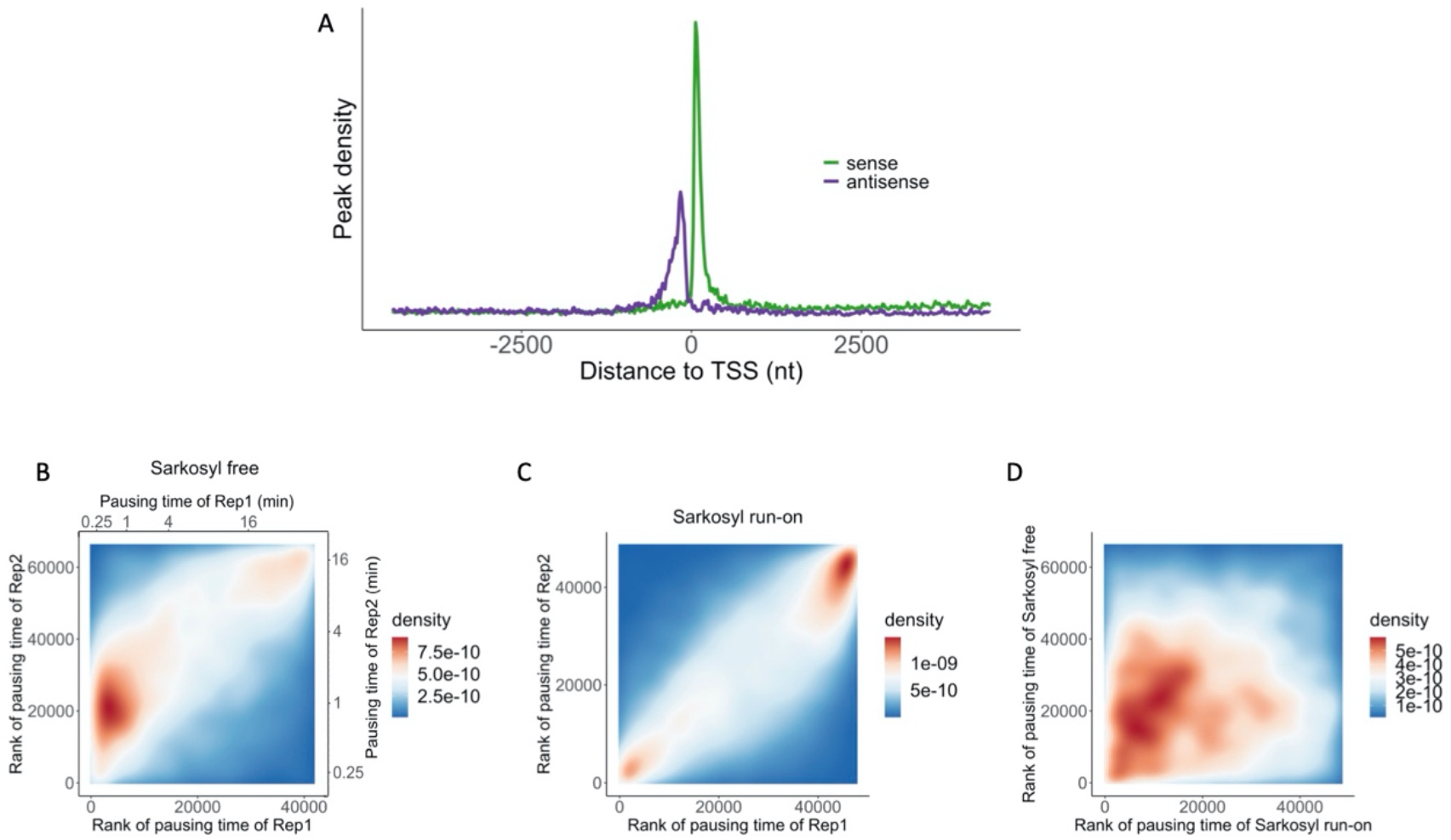
Consistency of TV-PRO-seq. (**A**) Distributions at TSS of sense and antisense reads of a replicate TV-PRO-seq sample confirms library qualities. (**B**) 2D density plot shows correlated pausing times between two replicates of TV-PRO-seq samples (sarkosyl free run-on). Top X axis and right Y axis refer to the pausing times of the two replicates. Bottom X axis and left Y axis refer to the ranks of pausing time. (**C**) 2D density plot shows correlated pausing times between two replicates of samples with sarkosyl in the run-on buffer. As the real pausing times cannot be estimated, only the ranks of pausing times are shown. (**D**) As (C), but comparing samples with or without sarkosyl, demonstrating a complex pattern.

**Fig. S3.**
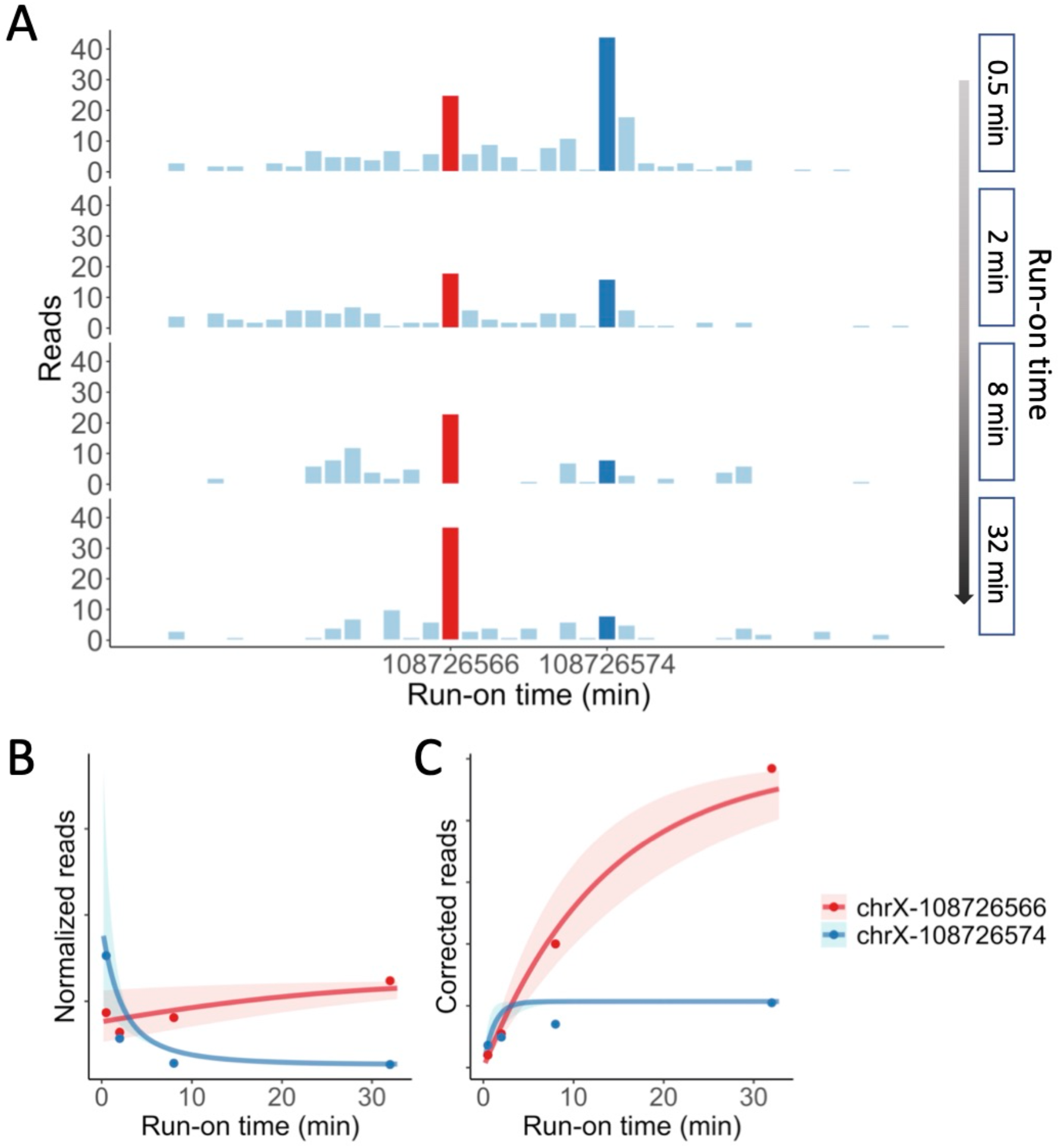
Example peaks for pausing time estimation of replicate. Read numbers normalised by total-genome reads, rescaled (by 10^7^), as used for model fitting. (**A**) Normalised reads of the example peaks of Figure 1C from the replicate data. (**B**) The resulting curve fits of peaks in (A). The shaded regions correspond to lower and upper quartiles of the posteriors. (**C**) Read numbers and saturation curves from the two example peaks as in Figure 1D, obtained from the replicate-experiment data.

**Fig. S4.**
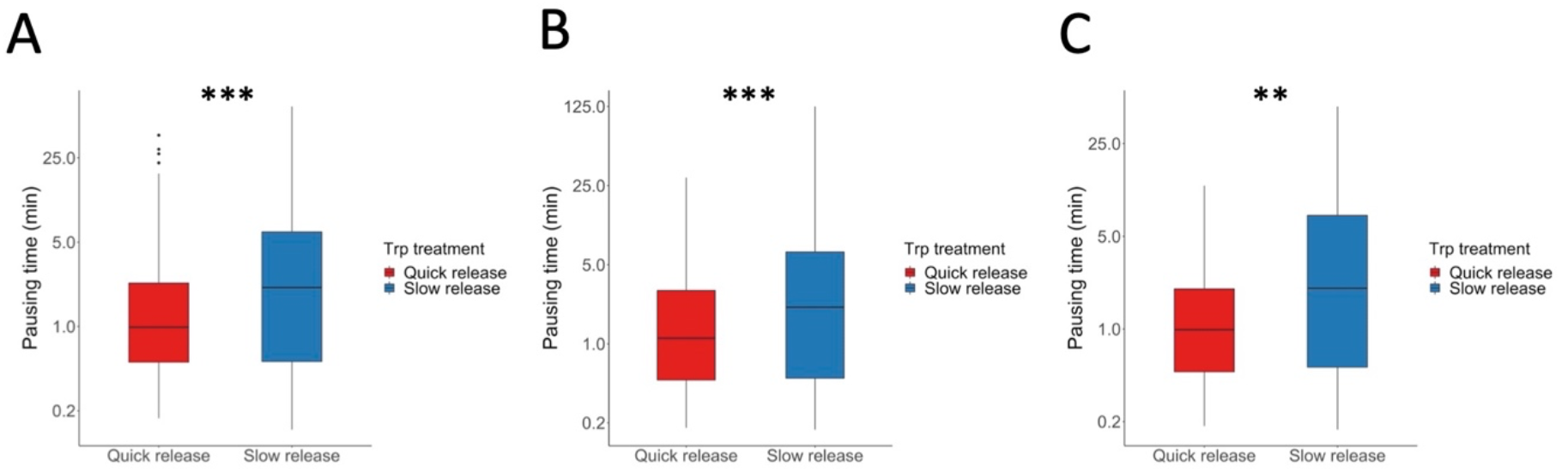
(A) 5,726 peaks were selected that had been identified as pausing sites in both TV-PRO-seq and Trp treatment following sequencing. Pausing peaks were grouped according to the foldchanges of polymerase occupancy after 10min Trp treatment. The top 20% of these were identified as ‘Quick release’, the bottom 20% as ‘Slow release’. P <10^-19^, Mann-Whitney U test. (B) As (A), for 1,107 peaks within 500bp downstream of TSS. P <0.001, Mann-Whitney U test. (**E**) As (A), for 547 peaks within 100bp downstream of TSS. P <0.01, Mann-Whitney U test.

**Fig. S5.**
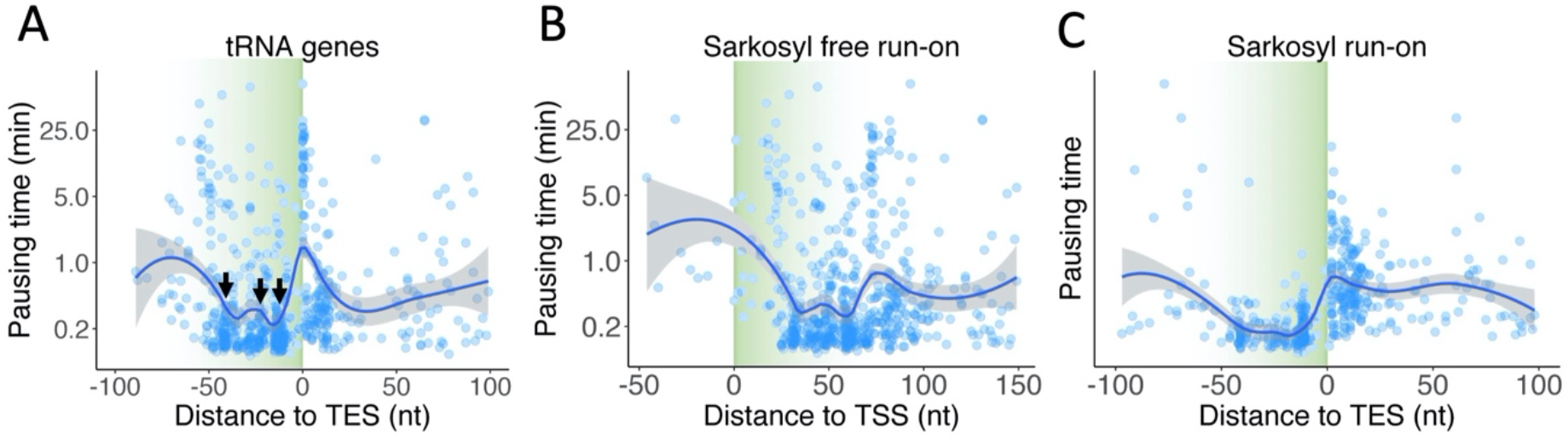
Pausing times and positions at tRNA genes. (**A**) Pausing times and positions at tRNA genes. Each dot corresponds to a pausing peak. The blue line corresponds to the moving average with the grey shading indicating its 0.95 confidence interval (LOESS fit). The metagene is aligned at the TES, where the pausing times interestingly increase. Selected common pausing sites at −42, −23 and −14 are marked with arrows. Due to the genes’ shortness, alignment at TSSs produces a similar plot (Fig. S4A). **(B)** As (A), pausing sites aligned with TSS. (**C**) As (A), under sarkosyl run-on conditions.

**Fig. S6.**
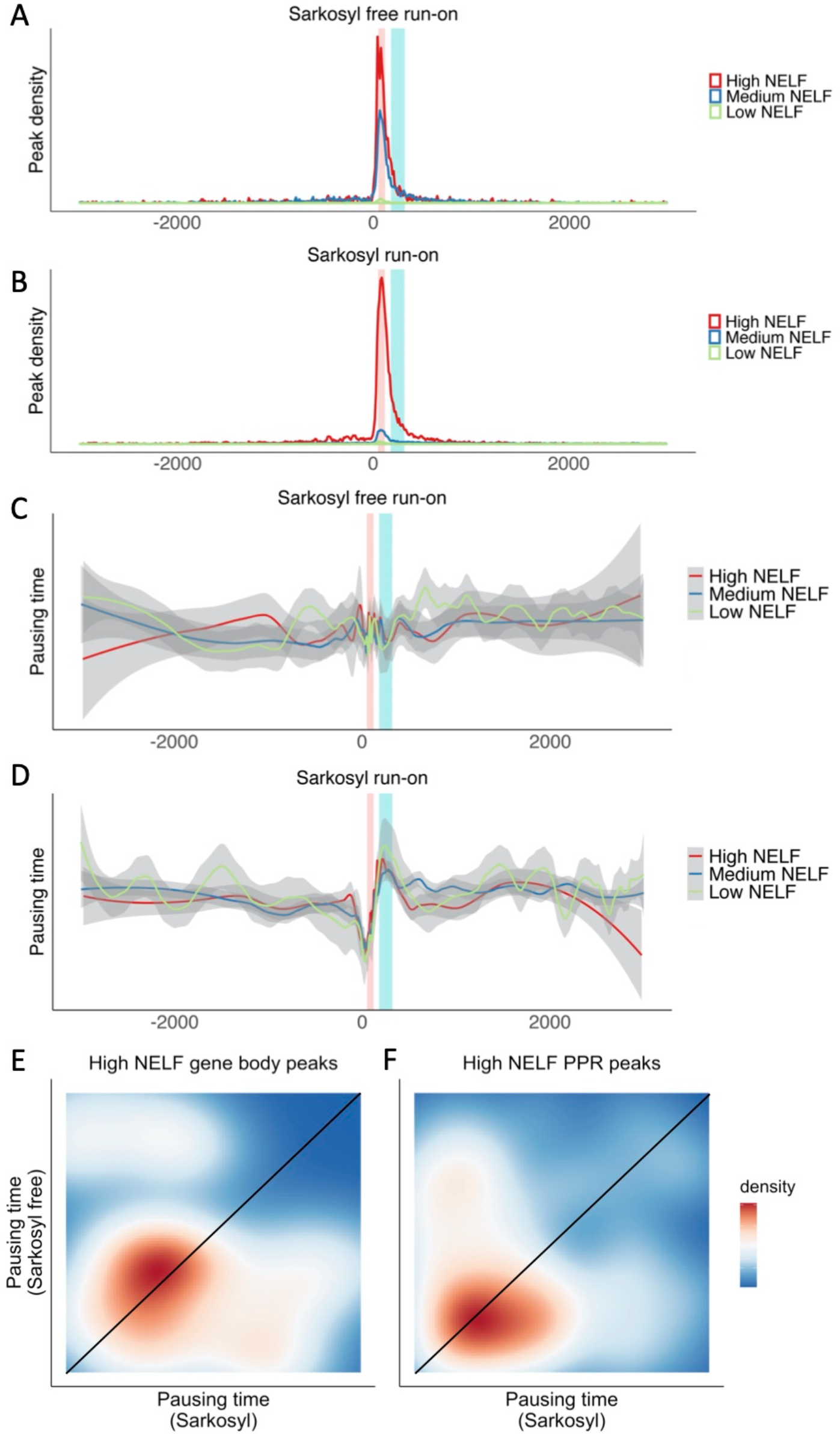
Pausing frequency and time of polymerase around TSS. (**A**) Density plot of peaks at different NELF levels within +/-3000nt of TSSs without sarkosyl during the run-on reactions. The red line represents high NELF signal within +/-80nt around peaks, blue medium and green low, respectively. (**B**) Similar to (A), for sample with sarkosyl run-on. (**C**) Moving average (LOESS fit) of pausing times of peaks with different NELF levels within +/-3000nt of TSSs in sarkosyl-free sample. The grey shading indicates the 0.95 confidence interval. Classification of peaks is the same as in (A). (**D**) Similar to (C), for sarkosyl run-on sample. (**E**) 2D density plots show the pausing time rank of the equivalent peaks in sarkosyl sample and sarkosyl-free sample. The black line reflects peaks with intermediate influence on pausing time by sarkosyl. Peaks above the black line correspond to pausing sites releasing paused polymerase after sarkosyl treatment. Only peaks after first 500bp of genes and have the top 10% NELF level have been selected. (**F**) Similar to (E), for peaks with in first 120bp and 10% NELF level.

**Fig. S7.**
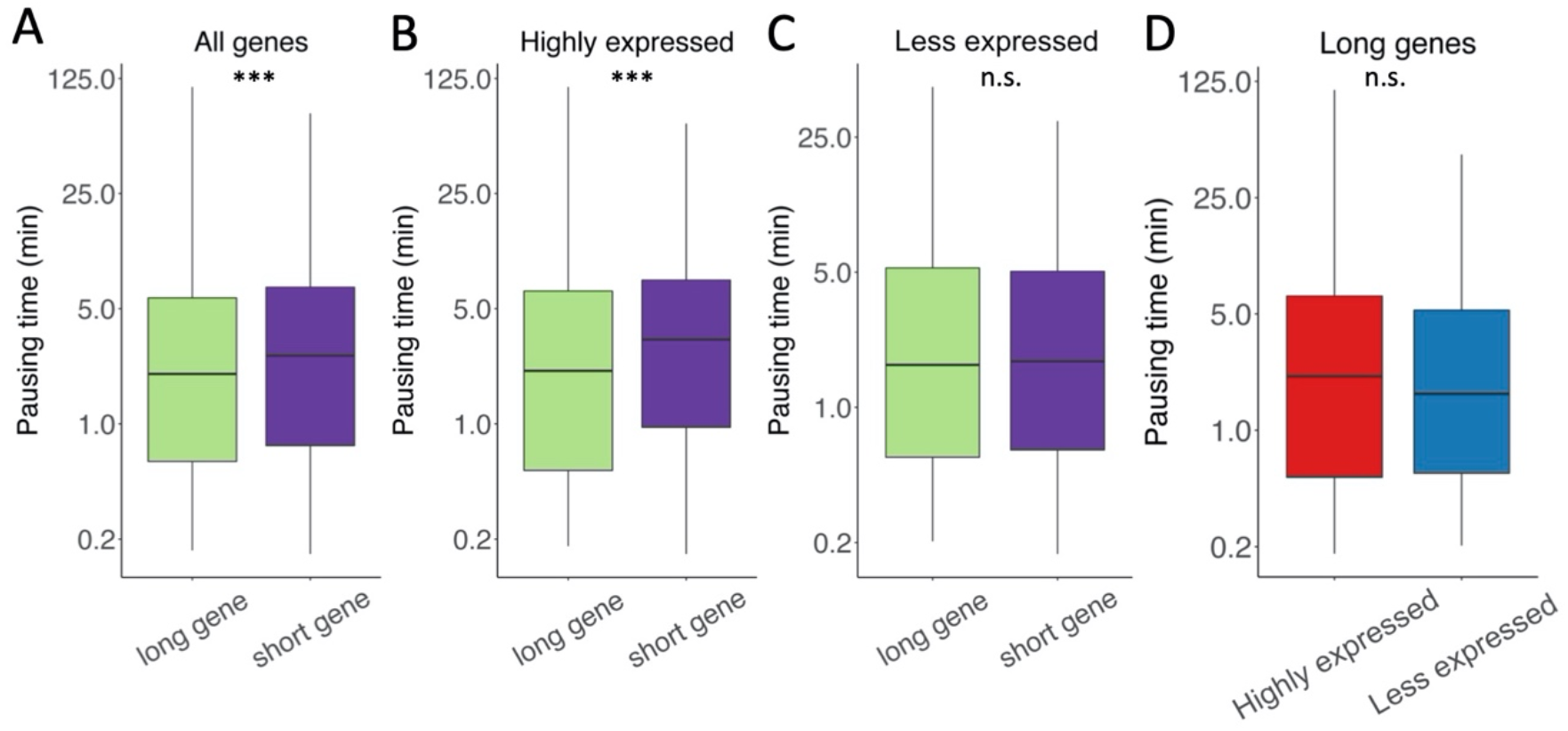
Influence of gene length on pausing times of genes with different expression levels. **(A)** Pausing times of peaks within long genes (>100kb, cyan) and short genes (3kb-10kb, purple), respectively. P< 10^-6^, Mann-Whitney U test. **(B)** Pausing times of peaks in highly expressed genes (top 20% expression). Group is similar to (A). P< 10^-4^, Mann-Whitney U test. **(C)** Pausing times of peaks in lowly expressed genes (bottom 20% expression). Group is similar to (A). Not significant, Mann-Whitney U test. **(D)** Pausing times of peaks from long genes (>100kb). Genes were further selected by expression level (highly expressed genes & less expressed genes, top & bottom 20% quantiles of expression levels, respectively).

**Fig. S8.**
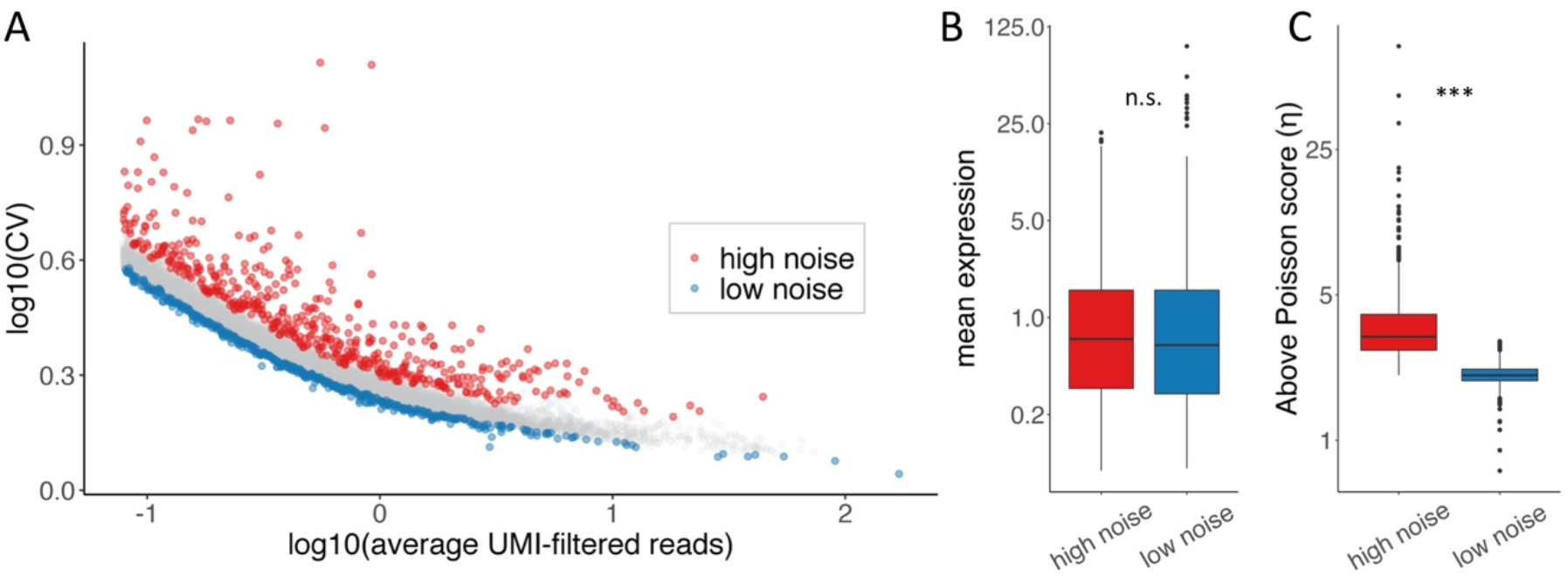
(**A)** Selection of high/low-noise genes. (**B**) Mean expression level does not show significant difference between high noise and low noise genes. (**C**) The above Poisson score *η* shows a significant difference between high noise and low noise genes (P < 10^-220^, Mann-Whitney U test), confirming the noise selection via an alternative noise measure.

**Fig. S9.**
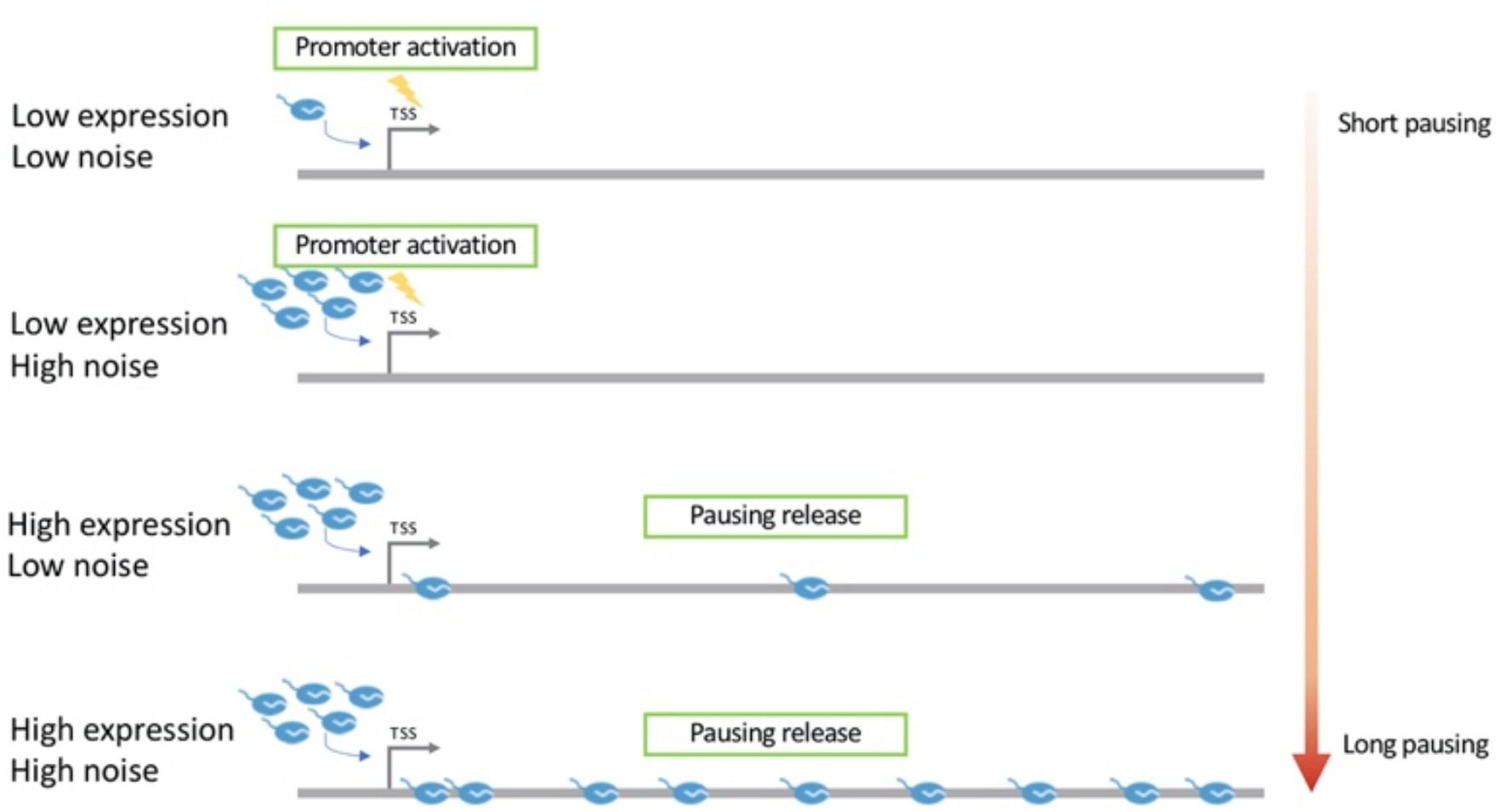
Schematic illustration of pausing profiles of different groups of genes. The rate-limiting step of lowly expressed genes is considered to be the promoter activation / transcription initiation. The number of polymerases initiated upon each promoter activation define its transcriptional noise (burst size). The promoters of highly expressed genes are always ready for initiation. Pausing therefore becomes the rate-limiting step for noisier genes. More and longer pausing allows for a higher polymerase load on the gene body which can generate a burst of transcription upon release.

**Fig. S10.**
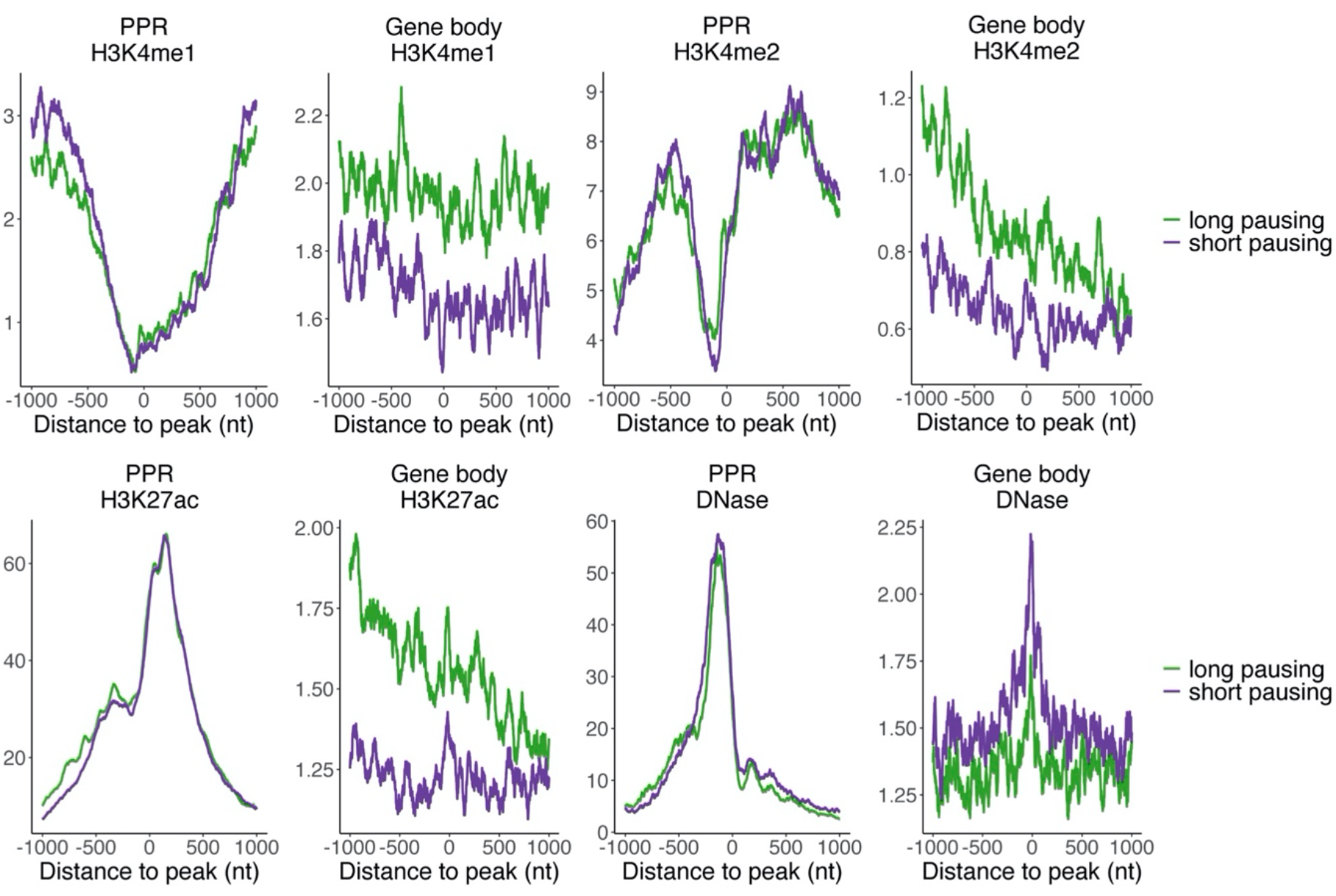
The average signal (AU) of H3K4me1, H3K4me2, H3K27ac and DNase-seq. ChIP-seq coverage around long/short pausing sites as determined by TV-PRO-seq.

**Fig. S11.**
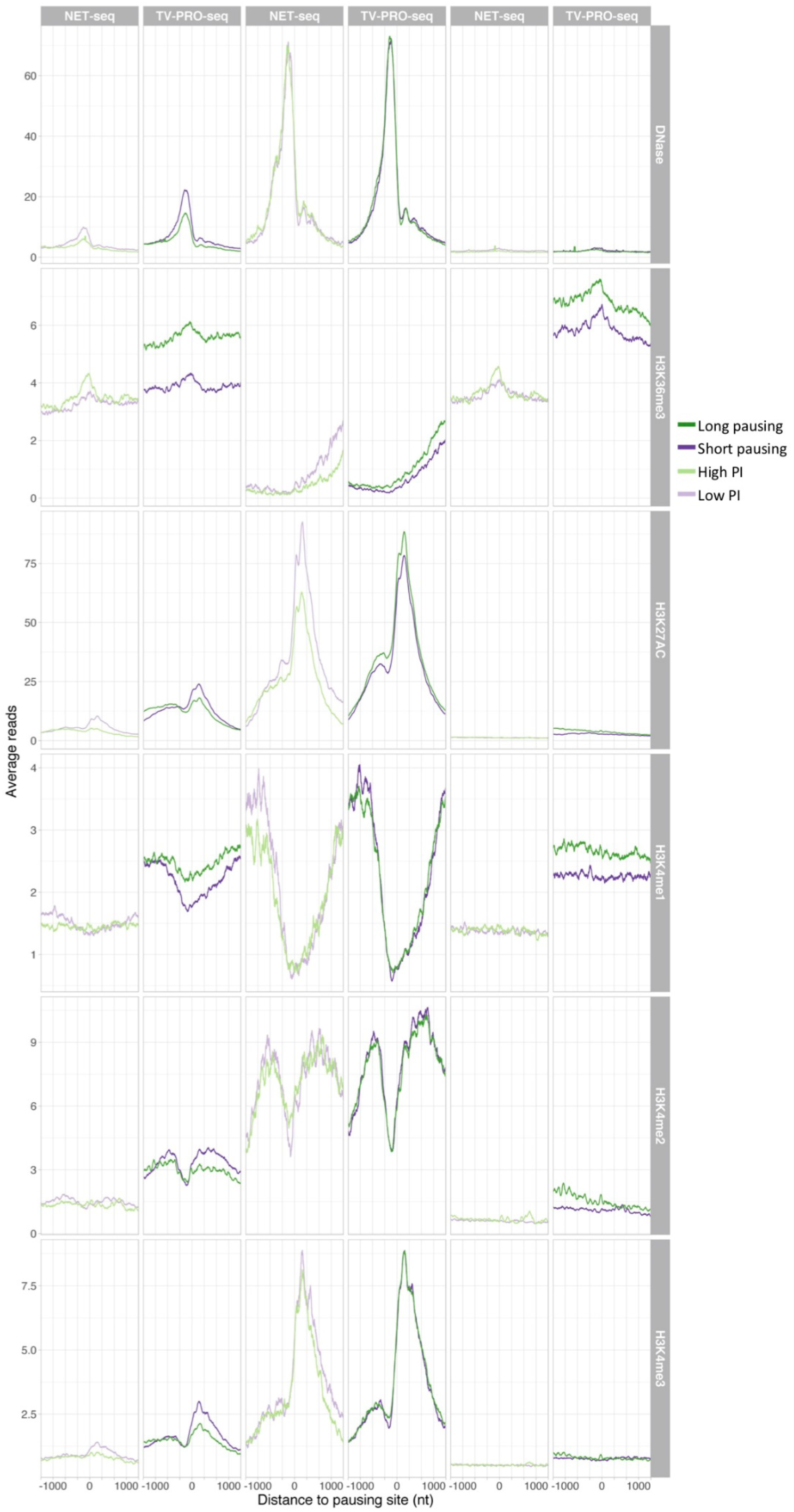
Comparison of chromatin state profiles for TV-PRO-seq and NET-seq/PI. Light purple and light green lines represent the low PI and high PI (Supp. Methods) pausing positions from NET-seq data, respectively. Dark purple and dark green represent the short pausing and long pausing positions from TV-PRO-seq data, respectively. The type of chromatin feature (as determined by DNase-seq or ChIP-seq) is shown on the right hand side; all peaks in the gene body, peaks in the region from TSS to +500 nt or TSS +2000 nt to TES - 1500 are shown in the first two, middle two and last two columns, respectively. The profiles are clearer for TV-PRO-seq in many cases, and often deviate from the NET-seq profiles, suggesting that TV-PRO-seq often produces better and sometimes different information.

## Footnotes

1 *b* is ideally estimated from the sample mean of read numbers at each of the 201 positions; however, many peaks are close to the TSS, which has many more reads downstream than upstream. To take account of this asymmetry, we assume that all the reads are downstream and average over the half-interval. This overestimates the background noise, and is thus a conservative estimate.

